# A gene silencing pipeline to interrogate human cDC1 and pDC development and functions

**DOI:** 10.1101/2023.05.16.540909

**Authors:** Xinlong Luo, Xavier Lahaye, Alix Jaeger, Paula Michea-Veloso, Nicolas Manel, Marc Dalod

## Abstract

Type 1 conventional dendritic cells (cDC1s) and plasmacytoid dendritic cells (pDCs) are thought to be critical for anti-tumor or antiviral immunity. In vitro differentiation systems have unlocked the ability to produce large numbers of these cells. However, a method is lacking to systematically identify the cell-intrinsic factors controlling their differentiation and functions that remain therefore poorly understood, in contrast to the situation in mice. Here, we developed a workflow for efficient gene silencing and its tracing in human cDC1s/pDCs generated in vitro. As proof of concept, we confirmed the key role of IRF8 in their development, and of IRF7/MyD88 in human pDC production of interferons-α/λ. We found that SAMHD1 and RAB7B promote human cDC1 differentiation, while SEPT3 promotes human pDC differentiation. We also found that PPT1 and RAB5 are required for optimal differentiation of pDCs and cDC1s. Finally, we identified BCL11A, PPT1 and RAB7 as novel HIV-1 restriction factors in cDC1s/pDCs. This approach will enable broader genetic screens to advance our understanding of human cDC1s/pDCs and harness them against viral infections or cancer.

## Introduction

The family of dendritic cells (DCs) regroups distinct types of mononuclear phagocytes that have in common a unique efficiency at activating naïve antigen-specific T cells upon their first encounter with their target antigen, a process coined T cell priming^1^. DCs encompass plasmacytoid dendritic cells (pDCs) and conventional dendritic cells (cDCs). pDCs are the main producer of type I (α/β) and III (λ) interferons (IFNs) upon the sensing of viral-type stimuli. cDCs are the most efficient cells for T cell priming and are further classified into cDC1s and cDC2s^2^. cDC1s excel in the priming of cytotoxic CD8 T cells, including via their unique efficacy at engulfing and processing antigens from dying cells, to present them in association with the class I major histocompatibility complex (MHC-I) molecules. This process, coined antigen cross-presentation, is crucial for immune defenses against cancer and intracellular pathogens^1^. cDC2s rather specialize in the recognition of, and defense against, extracellular parasites (e.g. worms), bacteria and fungi, via the priming of helper CD4 T cells and their functional polarization for the production of type 2 cytokines (helper type 2 or Th2 responses) or of interleukin-17 (Th17)^1^.

Most of the mechanistic knowledge we have on the functions of DCs and their molecular regulation has been acquired by using either mouse models^3^ or human DCs derived in vitro from monocytes (MoDCs)^4^. Human and mouse DCs populations share transcriptional signatures^5^. However, deterministic factors identified in mice do not necessarily play the same role in the human immune system and anti-microbial defense (e.g. TLR3^6^, IL23^7^ or IRF1^8^). MoDCs are a very valuable model that has led to major advances in our understanding of the molecular mechanisms underpinning key DC functions, including antigen processing and presentation, T cell priming and functional polarization. Yet, MoDCs strongly differ from cDCs on many accounts, including their overall molecular make-up^5, 9^, their adjuvant responsiveness^10^, their susceptibility to viral infections^11^ and their migratory behavior^12, 13^. Overall, MoDCs appear to be less efficient than cDC1s or even cDC2s or pDCs in promoting CD8 T cell responses against cancer, both in mice^14^ and in humans^15,16,17^. Hence, MoDCs are not an adequate surrogate model for many aspects of cDC biology.

The low frequency of cDCs and pDCs in all human tissues and their frailness has impeded their study ex vivo as well as their use in the clinic for adoptive cell therapy, in particular for cDC1s. Human DC immunodeficiencies provide a forward-genetic-based approach to deciphering human DC biology^18^. However, it remains necessary to implement reverse genetics-based approaches to mechanistically dissect human DC biology. To dissect the molecular mechanisms controlling pDC activation, many teams have been using tumor cell lines derived from blastic plasmacytoid dendritic cell neoplasms (BPDCN), such as GEN2.2^19^ or CAL1^20^. However, BDPCN cells have undergone major genetic perturbations, including multiple mutations in epigenetic modifiers^21, 22^. This raises questions on the extent to which the study of the functions BDPCN cell lines and their molecular regulation is relevant to normal primary pDCs or may be affected by malignant transformation^23^. Indeed, BPDCN cell lines differ from primary pDCs on different accounts, including a lesser ability to produce IFNs in response to viral-type stimuli, and an abnormal dependency for this function on the transcription factor interferon regulatory factor 5 (IRF5)^24, 25^, instead of IRF7 as used in primary pDCs^26, 27^. Thus, there is a strong need for protocols enabling the generation and manipulation of human pDCs and cDC1s in vitro under conditions where these cells constitute faithful models of their in vivo counterparts but are as amenable to experimental manipulation as MoDCs or BPDCN cell lines.

CRISPR/Cas9 constitutes an emerging approach to study gene functions in human DCs. To this date, this has been limited to the study of MoDCs, due to the impact on cell yield of CRISPR/Cas9 ribonucleoprotein electroporation^28, 29^. In addition, bulk knock-out (KO) of DC precursors generates a mixture of WT and KO cells. However, because DC differentiation and functions are profoundly affected by cell-cell interaction and cytokines, such as TNF or type I interferons, a method to trace gene-deficient DCs is essential to parse out cell-intrinsic gene function from secondary effects of cell-extrinsic activities. Tracing of KO cells using CRISPR/Cas9 requires the concomitant knock-in of a report cassette, which creates additional challenges.

Different methods have been reported to generate in vitro human pDCs, cDC1s or cDC2s^30^, including a cost-effective, highly efficient and feeder layer-dependent protocol for in vitro differentiation of human cord blood (CB) CD34^+^ hematopoietic stem cells (HSCs) into bona fide cDC1 and pDCs that our team developed recently^31^. However, likewise to their in vivo counterparts, these in vitro derived cDC1s and pDCs are highly resistant to viral infection or transduction^11^, making very challenging their genetic manipulation for deciphering the molecular mechanisms regulating their development and functions. Therefore, we aimed at adapting our pipeline for cDC1/pDC differentiation by transducing the expanded HSCs prior to inducing their differentiation into cDC1s and pDCs, instead of transducing the already differentiated cDC1s and pDCs. We optimized different parameters to obtain a high transduction rate of both the expanded HSCs and their cDC1/pDC progeny, without any strong perturbation of the expansion and differentiation of the cells, and no spontaneous activation of the obtained cDC1s/pDCs, when using control lentivectors. Importantly, the lentivirus-based approach enables the tracing of the modified cells that is required to ensure the cell-intrinsic nature of gene activities. We then validated this pipeline by recapitulating upon gene knock-down of *IRF8* or *IRF7* in vitro the impact on the development or adjuvant responsiveness of cDC1s/pDCs that had been previously identified in patients suffering from the corresponding primary immune deficiencies^26, 32^.

Using our pipeline, we performed a small-scale screen by knocking-down 10 candidate genes and assessing whether it impacted cDC1 or pDC biology, including (1) their development, (2) their ability to produce innate cytokines in response to Toll-like receptor triggering, and (3) their sensitivity to infection by type 1 human immunodeficiency virus (HIV-1).

We choose to study the three transcription factors ID2, DC-SCRIPT and BCL11A, because, in mice, the first two have been shown to promote cDC1 development^33,34,35,36^ and the last the development of pDCs^37, 38^. We were able to extend these findings to human DCs, for the first time to the best of our knowledge.

We also screened five small GTPases, RAB7B, RAB15 and SEPT3 because they are selectively expressed at high level in pDCs or cDC1s^5, 11, 39^, but their functions in these cells is largely unknown, as well as RAB5A and RAB7 for comparison. Rab small GTPases perform key functions in intracellular vesicle trafficking in the endocytic, exocytic, and transcytic pathways^40^. Some Rab family members control antigen presentation by mononuclear phagocytes or their adjuvant responsiveness, by modulating intracellular routing and degradation of endocytosed antigens (RAB7^41^ and RAB43^42^), MHC-I molecules (RAB11A^43^), or pattern recognition receptors (RAB7B^44, 45^ or RAB11A^46, 47^). Rab small GTPases can also impact pathogen entry/egress, including in the late phase of HIV-1 life cycle where active membrane trafficking processes are involved in macrophages^48, 49^ and T cells^50^. For example, RAB7A promotes HIV-1 envelop maturation and particle release by facilitating Vpu-induced degradation of host restriction factor BST2 and helping the cleavage process of the envelop glycoprotein for optimal infectivity of the progeny virus^51^. We have previously reported that RAB15 contributes to cDC1 resistance to HIV-1 infection by restricting membrane fusion at early phase of the viral life cycle^11^. Here, we extended this study to pDCs, since they also express high levels of RAB15. Septins are small cytoskeleton-binding GTPases forming heteromers that instruct the specific intracellular localization of interacting proteins and can also define membrane compartments by defining diffusion barriers, from yeast to men^52^. Septins associate with syntaxins, a family of proteins directly involved in phagocytosis^53^. In addition, Septins can generate cage-like structures that entrap intracytosolic bacteria^54^ and limit their replication by delivering them to lyzosomes or autophagosomes^55, 56^. Yet, the function of SEPT3 in cDC1s is unknown.

We also screened two candidate viral restriction factors in addition to RAB15: palmitoyl-protein thioesterase 1 (PPT1) since it has been shown to restrict infection of mouse cDC1s by vesicular stomatitis virus^57^, and SAMHD1 considering its well-known role in MoDCs^58,59,60,61,62^. Indeed, the fact that cDC1 sensitivity to HIV-1 infection was only slightly increased upon RAB15 knock-down^11^ suggested that cDC1 were protected by other restriction factors.

Our novel pipeline for gene silencing during the in vitro differentiation of HSCs into bona fide human cDC1s and pDCs enabled us to (1) extend to humans the demonstration previously achieved in mice that the transcription factors ID2 and DC-SCRIPT versus BCL11A promote cDC1 versus pDC development respectively, (2) unravel a novel role of SAMHD1 in balancing HSC differentiation into cDC1s versus pDCs, (3) identify small GTPases that control the development or activation of human cDC1s or pDCs, (4) extend to pDCs the restriction factor activity of RAB15 previously discovered in cDC1s, and demonstrate that RAB7 and BCL11A also limit HIV-1 replication in cDC1s and pDCs.

## Results

### Pipeline development to genetically edit human cDC1s/pDCs

We confirmed that differentiated cDC1s and pDCs were largely refractory to lentiviral transduction, even when using concentrated lentivectors (**Extended Data Fig. 1A**). To overcome this technical bottleneck, we optimized our protocol to achieve a high transduction rate of CB HSCs, with maintenance of a high proportion of transduced cells in their differentiated pDC and cDC1 progeny (**Extended Data Fig. 1B-G**; **Fig. 1A-B**). We first tested modifications of the medium and cytokines used for the culture of the HSCs, to select conditions promoting a higher transduction efficacy. Using X-VIVO 15 serum-free medium without supplements, and the cytokine/growth factor cocktail FLT3L/SCF/TPO (FST), provided a better transduction rate than the four other culture conditions tested (**Extended Data Fig. 1B**). The transduction rate of the HSCs was further increased by optimizing five additional parameters (**Extended Data Fig. 1C**): (i) using higher molarities of infection (MOI) upon increasing lentivector titers by concentration, (ii) promoting cell-to-cell contacts upon using 96-well U-bottom plates instead of 24-well plates, and hence also (iii) performing the transduction in a small volume, (iv) replacing protamine with poloxamer 407 that had been reported to promote higher transduction rates of HSCs^63^, and (v) adding to our culture protocol a “priming phase” by culturing expanded HSCs in the transduction medium without poloxamer (“priming medium”) for one day prior to their exposure to lentivectors. This optimized protocol yielded higher percentages of transduced HSCs (up to 90%) and also promoted a better cell survival, as compared to the semi-optimized protocol (**Extended Data Fig. 1D**). After two to three weeks of differentiation, the percentages of the transduced cells were maintained at high levels, similar to those of their parental HSCs that had been transduced with the optimized protocol (**Extended Data Fig. 1E**). Seeding more HSCs on the feeder layer increased their production of cDC1s during the differentiation (**Extended Data Fig. 1F**). Hence, we selected to seed 2×10^4^ HSCs/ml for the differentiation phase, as the best compromise between the cost of the HSCs and the increase in their differentiation into cDC1s. The final scheme of the optimized pipeline is illustrated in **Extended Data Fig. 1G**. Under these experimental conditions, when using BFP-expressing shRNA lentivectors, both transduced and non-transduced cDC1s/pDCs were clearly detectable by flow cytometry in the same culture well (**Fig. 1B**). Thus, it allowed comparing the gene-modified versus control cells side-by-side, to assess the cell-intrinsic effects of each genetic manipulation in the DC lineages as compared to other cell types, even in the face of eventual indirect effects changing the overall differentiation conditions in the well. Indeed, to rigorously assess how the genetic manipulation under study specifically affected the development of cDC1s or pDCs in a cell-intrinsic manner, we defined a parameter named “differentiation efficacy” that assessed the increase or decrease in the proportion of transduced cells within the cDC1 or pDC populations, normalized to a similar ratio calculated for the whole population of viable cells in the same well (**Fig. 1C**).

**Fig. 1.**
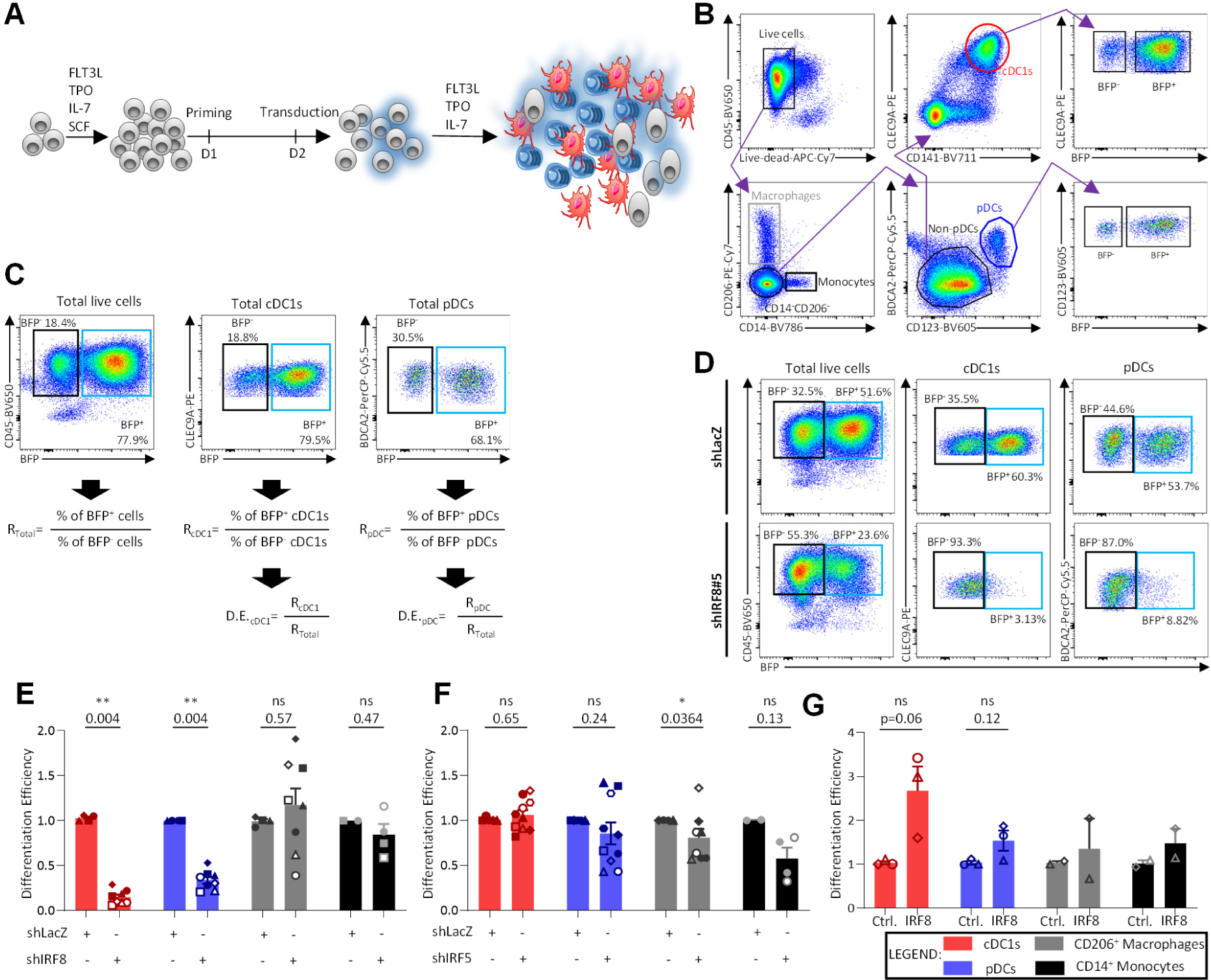
Optimization and validation of a workflow to produce gene-modified primary human cDC1s and pDCs. **A)** Schematic illustration of the overall workflow. **B)** Gating strategy for identification of transduced and untransduced cDC1s and pDCs, as well as CD14^+^ monocytes and CD206^+^ macrophages. **C)** Definition of “Differentiation Efficacy”, illustrated with samples transduced by a BFP-reporter lentivector. **D-G)** Impact of IRF8 and IRF5 knock-down on cDC1 and pDC development. **D)** FACS plots from representative cultures of HSCs transduced with control (shLacZ) or shIRF8 lentivectors. **E-F**) Impact of IRF8 (**E**) and IRF5 (**F**) on the differentiation efficacy of cDC1s, pDCs, monocytes and macrophages. The results are shown from four independent experiments, with four CB donors (symbol shapes), and two different shRNA (full versus empty symbols) for IRF8; five independent experiments, with five CB donors (symbol shapes), and two different shRNA (full versus empty symbols) for IRF5. **G**) Impact of IRF8 overexpression on cDC1 and pDC development. HSCs were transduced with a lentivector encoding human IRF8. Their output was assessed after 18 days in differentiation culture, according to the pipeline shown in panel A. The results are shown from three independent experiments, with three CB donors. For panels (**E-F**), statistical analyses were performed using a two-tailed non-parametric Mann-Whitney test. For panel (**G**), statistical analyses were performed using a paired two-tailed t test. *, p<0.05; **, p<0.01; ***, p<0.001; ****, p<0.0001; ns, not significant.

**Extended Data Fig. 1.**
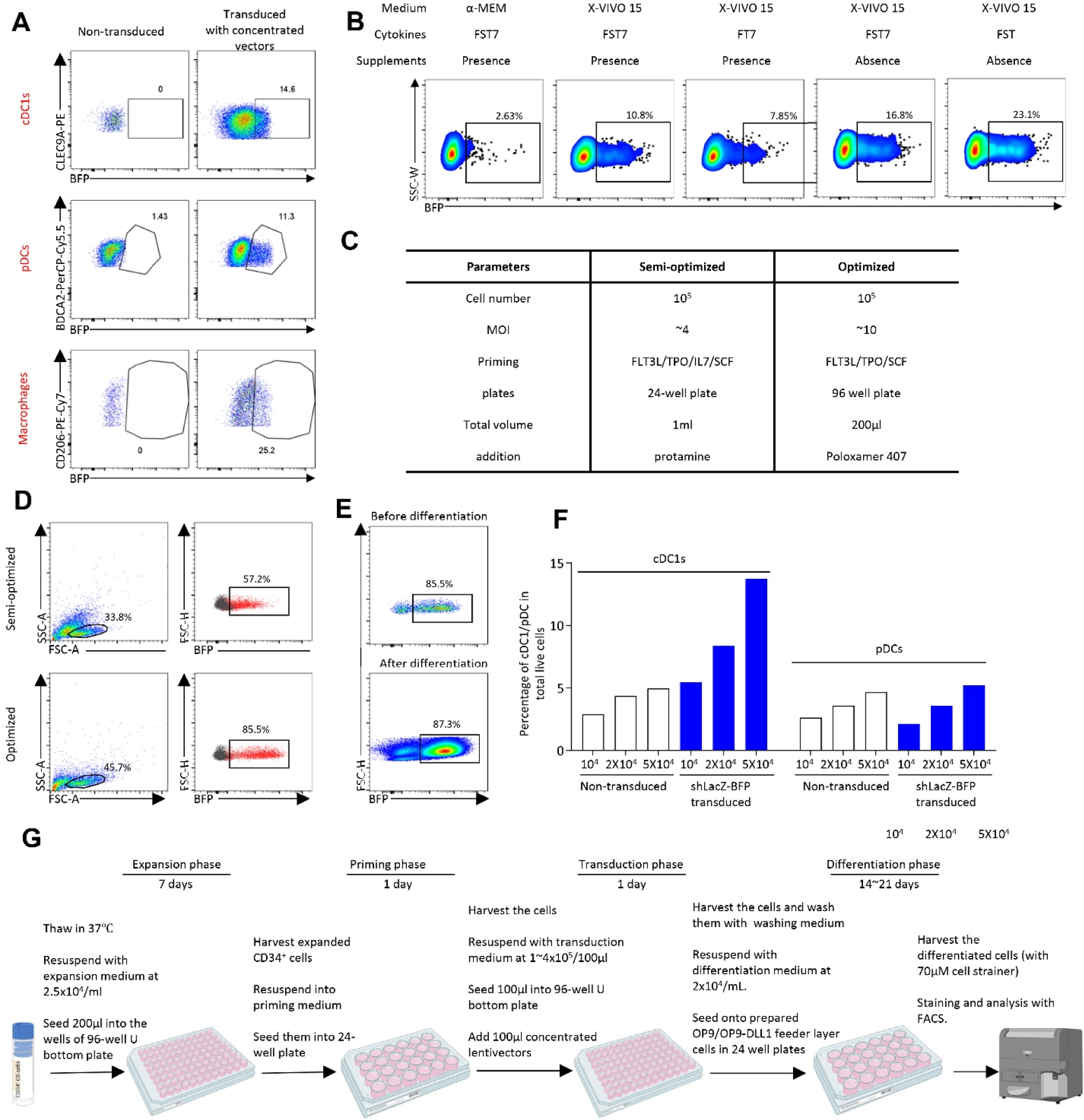
Parameters optimized for high efficacy transduction of primary human cDC1s and pDCs. **A)** FACS plot showing the transduction rate of cDC1s, pDCs and macrophages with concentrated lentivectors. **B)** FACS plot showing the effects of medium, cytokine selection and supplement presence on the transduction rate of expanded CD34^+^ cells. **C)** List of the parameters optimized to increase the transduction rate of progenitors. **D)** FACS plot showing the transduction rate of progenitors comparatively between the semi-optimized and optimized protocols. **E)** FACS plot showing the proportion of transduced cells before and after the differentiation process, with the optimized protocol. **F)** Impact of the initial number of cells seeded on the generation of cDC1s and pDCs. **G)** Final detailed scheme of the optimized pipeline.

### Role of IRF8 in human cDC1 and pDC development

We aimed at establishing a proof of concept to validate the suitability of our protocol to assess the impact of the knock-down of candidate genes in human HSCs on their ability to differentiate into cDC1s or pDCs. To this aim, we targeted IRF8, because this transcription factor has been demonstrated to be pivotal for cDC1 and pDC development both in mice as assessed with mutant animal models^33, 64,65,66,67^ and in humans as observed in patients with loss-of-function alleles^32, 68,69,70^. We used two different shRNAs that we had validated to strongly decrease IRF8 protein expression in THP-1 cells (**Fig. S1A**). IRF8 inactivation strongly and specifically impaired HSC differentiation into both cDC1s and pDCs (**Fig. 1D-E**). The differentiation into CD206^+^ macrophages and CD14^+^ monocytes was not affected (**Fig. 1E**). In contrast, IRF5 inactivation reduced HSC differentiation into monocytes with a similar trend for macrophages but no change for cDC1s and pDCs (**Fig. 1F**). Conversely to its inactivation, IRF8 overexpression tended to increase the differentiation of HSCs into cDC1s (**Fig. 1G**). Thus, these results demonstrate that our culture system recapitulates in vitro the key requirement of IRF8 but not IRF5 for human cDC1 and pDC development. Thus, for the first time to our knowledge, we have achieved combining genetic manipulation of HSCs with an in vitro system recapitulating their physiological differentiation into human cDC1s and pDCs, enabling to identify the underlying molecular mechanisms.

### Role of IRFs and adaptor molecules in pDC/cDC1 activation

Next, we harnessed our pipeline to investigate the role of IRF7 versus IRF5 and IRF8 in primary human pDCs derived in vitro from HSCs. We selected three different shRNAs significantly diminishing IRF7 protein expression in transduced pDCs (**Fig. S1B-C**). IRF7 inactivation strongly decreased pDC production of IFN-α and IFN-λ, but not TNF, in response to stimulation with the Toll-like receptor (TLR)7 ligand R848 (**Fig. 2A-B**), whereas this was the case neither for IRF5 (**Fig. 2C**) nor for IRF8 (**Fig. 2D**). Thus, primary human pDCs generated in vitro critically require IRF7 for IFN-I/III production in response to viral-type stimuli, whereas IRF5 is dispensable for this function, as observed for primary blood pDCs^26, 27^, in contrast to the opposite results observed with BDPCN cell lines^24, 25^. Hence, results obtained with BDPCN cell lines should be considered with caution. Our pipeline will allow checking whether they hold in bona fide primary pDCs derived in vitro from HSCs and amenable to genetic manipulation.

**Fig. 2.**
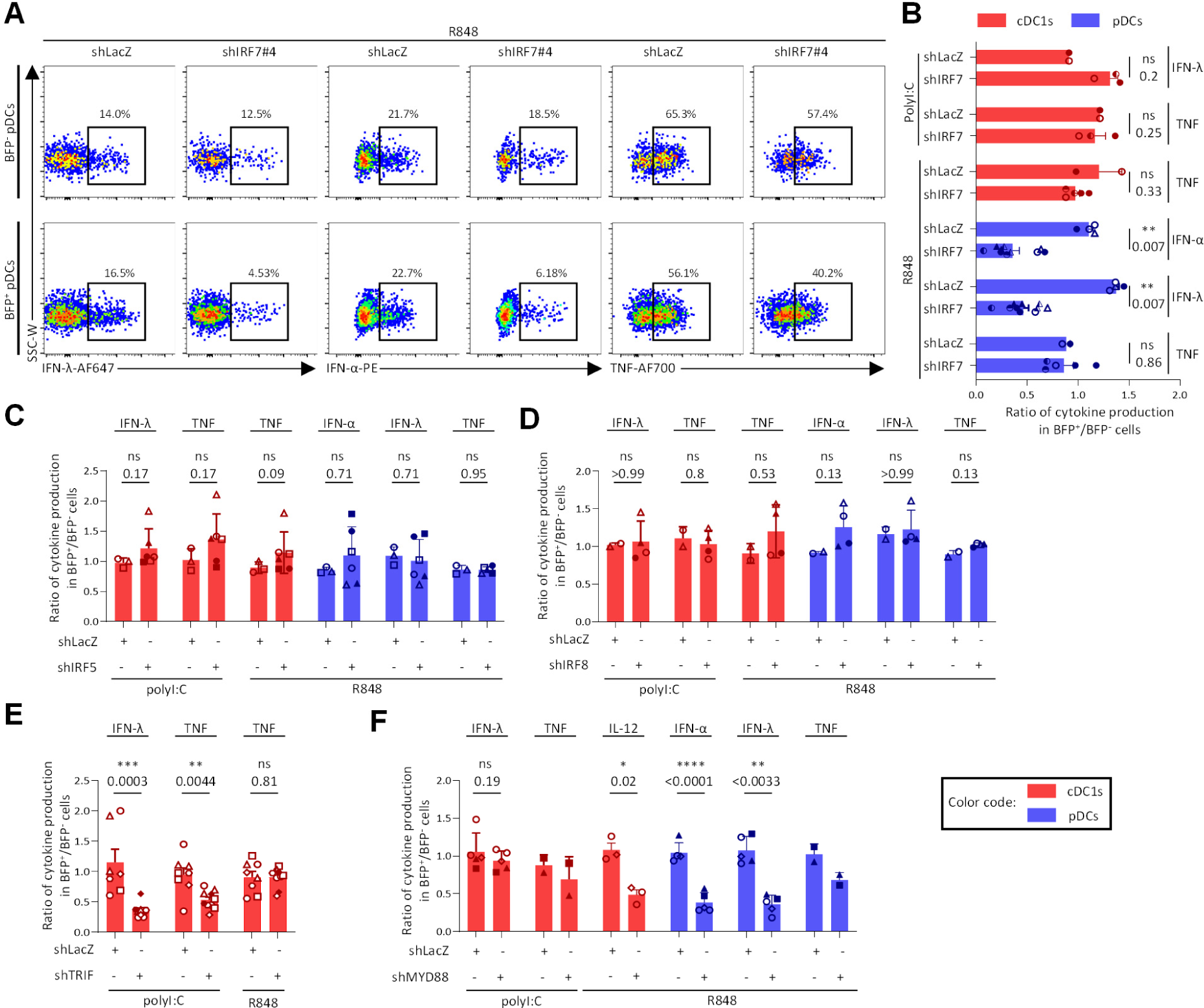
Role of IRF transcription factors and adaptor molecules in pDC and cDC1 cytokine responses to TLR triggering. Impact of the knock-down of IRF7 **(A-B)**, IRF5 (**C**), IRF8 (**D**), TRIF (**E**) and MYD88 (**F**) on cytokine production by cDC1 and pDC upon stimulation with TLR ligands. **A**) Representative FACS plots showing cytokine expression in pDCs from cultures of HSCs transduced with control (shLacZ) or shIRF7 lentivectors, upon stimulation with R848. **B-D**) Impact of IRF7 (**B**), IRF5 (**C**) and IRF8 (**D**) knock-down on cytokine production by pDCs or cDC1s. **E-F**) impact of TRIF (**E**) or MYD88 (**F**) knock-down on cytokine production by cDC1s (**E, F**) or pDCs (**F**). The results are shown from three independent experiments, with two distinct CB donors, and three different shRNA for IRF7; three independent experiments, with three distinct CB donors, and two different shRNA for IRF5; two independent experiments, with two distinct CB donors, and two different shRNA for IRF8; seven independent experiments, with five distinct CB donors, and two different shRNA for TRIF; and five independent experiments, with four distinct CB donors, and two different shRNA for MYD88. For panels (**B**, **C**, **D** and **E**), statistical analyses were performed using a two-tailed non-parametric Mann-Whitney test. For panel **F**, statistical analyses were performed using a two-tailed paired t test. *, p<0.05; **, p<0.01; ***, p<0.001; ****, p<0.0001; ns, not significant.

Distinct adaptor molecules promote responses to different TLRs. In mice, MyD88 is required for responses to TLR7, TLR9 and TLR8, including for pDC IFN-I/III production in response to viral-type stimuli ^71, 72^, whereas TRIF is required for TLR3 response, including the unique ability of cDC1s to produce high levels of IFN-III in response to stimulation with polyI:C^73^. Hence, we used our pipeline to determine which adaptor molecule was required for cytokine production by human pDCs or cDC1s in response to TLR triggering (**Fig. 2E-F, Fig. S1D-G**). As expected, cDC1 IFN-λ and TNF expression in response to polyI:C stimulation was significantly reduced upon TRIF knock-down (**Fig. 2E**), whereas this was not the case for cDC1 TNF production in response to R848 that is known to trigger TLR8 in cDC1s (**Fig. 2E**). Conversely, in response to R848, cDC1 production of IL-12 and pDC production of IFN-α and IFN-λ were decreased by MYD88 knock-down (**Fig. 2F**). Thus, these results demonstrate that our culture system recapitulates in vitro the key requirement of TRIF versus MYD88 for TLR3-versus TLR7/8-triggered cytokine responses of human cDC1s and pDCs, and they demonstrate the suitability of our pipeline for gene edition via knock-down in human HSCs in order to study the molecular regulation of the responses of human cDC1s and pDCs to TLR stimulations.

### Role in human DCs of transcription factors identified in mice

The lack of adequate experimental models has prevented the extension to human DC types of the analysis of the role of transcription factors found to be essential for the differentiation of mouse cDC1s or pDCs. By using our pipeline with validated shRNA lentivectors (**Fig. S2**), we showed here that, likewise to what had been reported in mice^33,34,35,36^, both ID2 (**Fig. 3A)** and DC-SCRIPT (**Fig. 3B)** promoted human cDC1 development in vitro from HSCs. ID2 knock-down tended to increase the output of pDCs in some experiments (**Fig. 3A**), consistent with the demonstration in mice that ID2 favor cDC1 over pDC differentiation by inhibiting the expression of the master transcription factor instructing pDC fate, TCF4^74^. The cDC1s knocked-down for ID2 harbored a significant but slight decrease in TNF production upon polyI:C stimulation (**Fig. 3C)**, whereas, on the contrary, the cDC1s knocked-down for DC-SCRIPT tended to harbor a slight increase in TNF production upon polyI:C or R848 stimulation (**Fig. 3D)**. Unexpectedly, ID2 knock-down significantly reduced pDC IFN-λ production in response to R848, with a tenecny towards a reduction of TNF production also (**Fig. 3C**). BCL11A promoted both pDC development (**Fig. 3E**) and their production of IFN-α/λ in response to TLR7 triggering (**Fig. 3F**). BCL11A knock-down also tended to reduce the output of cDC1s (**Fig. 3E**). This result is consistent with those previously reported upon in vitro differentiation of mouse BCL11A^-/-^ fetal liver cells^37^, even though BCL11A^-/-^ mice harbor impaired development of pDCs but not of cDC1s^37, 38^. Thus, these results enabled us to demonstrate critical roles of ID2 and DC-SCRIPT for promoting human cDC1 development, and of BCL11A for promoting human pDC development, for the first time to our knowledge, which is an important translation to humans of previous knowledge discovered by using mutant mouse models.

**Fig. 3.**
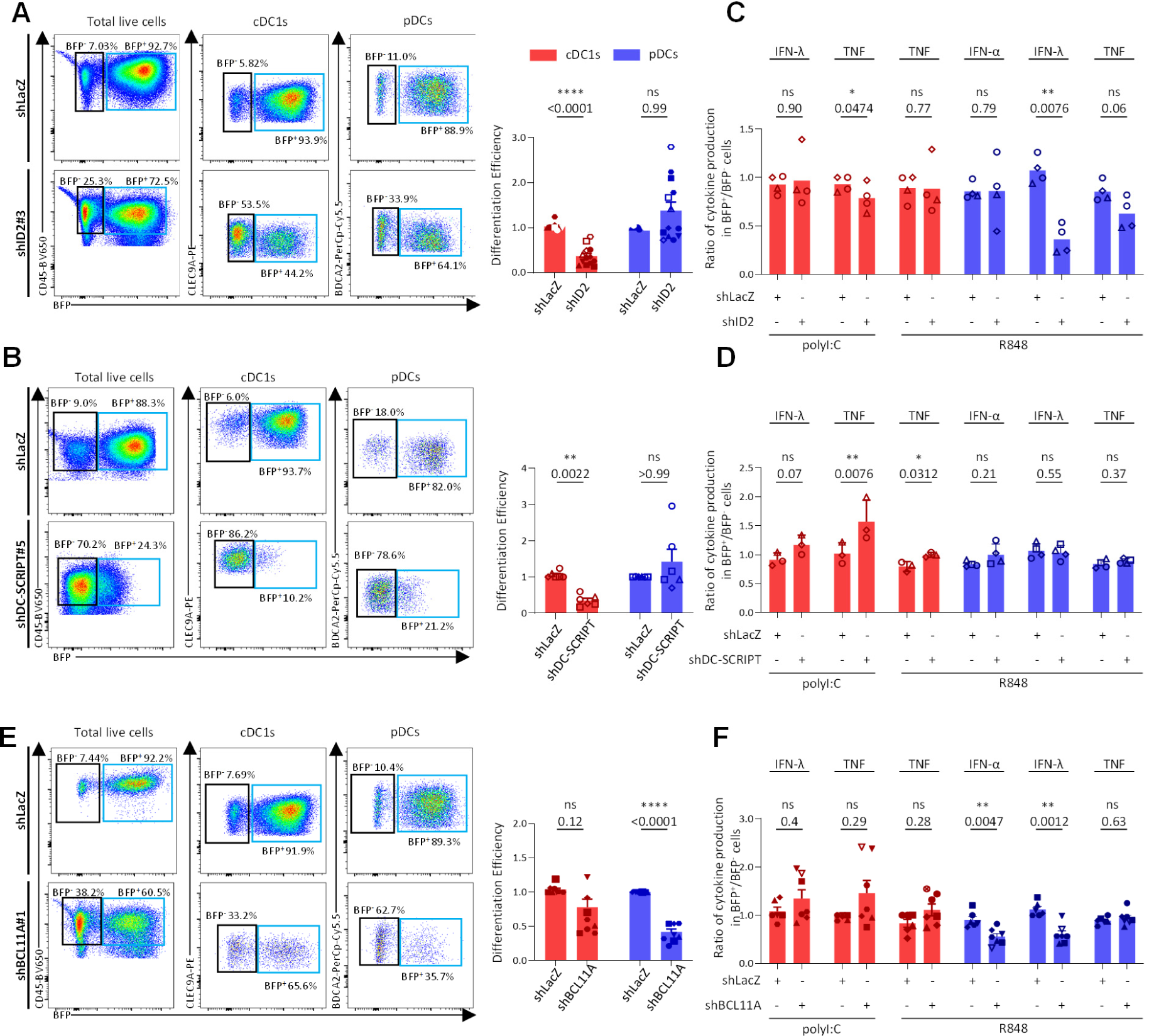
Impact of candidate transcription factors on the development and activation of human pDCs and cDC1s. Impact of the knock-down of *ID2* (**A, C**), *DC-SCRIPT* (**B, D**) and *BCL11A* (**E, F**), on human cDC1 and pDC development (**A**, **B**, **E**) or activation in response to TLR ligand stimulation (**C**, **D**, **F**). (**A**, **B**, **E**) for ID2, the results are shown from seven independent experiments, with five distinct CB donors and two different shRNA (full versus empty symbols); for BCL11A: ten independent experiments, with six CB donors and one shRNA; for DC-SCRIPT: four independent experiments, with four CB donors and one shRNA. (**C**, **D**, **F**) for ID2, the results are shown from six independent experiments, with four distinct CB donors and one shRNA; for BCL11A: six independent experiments, with six CB donors and two shRNAs; for DC-SCRIPT: four independent experiments, with three CB donors and one shRNA. For panels **A**, **B**, **E** and **F**, statistical analyses were performed using a two-tailed non-parametric Mann-Whitney test. For panel **C** and **D**, statistical analyses were performed using a paired two-tailed t test. *, p<0.05; **, p<0.01; ***, p<0.001; **** p<0.0001; ns, not significant.

### Identification of small GTPases controlling human DC biology

We next screened five small GTPases, RAB7B, RAB15 and SEPT3 because they are selectively expressed at high level in pDCs or cDC1s^5, 11, 39^, but their functions in these cells is largely unknown, as well as RAB5A and RAB7 for comparison. We selected between 1 and 3 shRNA per target, based on their knock-down efficiency as assessed by western blot (**Fig. S3**). The knock-down of *RAB7B* (**Fig. 4A**) or *RAB5A* (**Fig. 4B**) strongly reduced the cDC1 output of the cultures, while slightly increasing or decreasing the pDC output, respectively. The knock-down of *RAB7B* also slightly increased cDC1 production of TNF upon R848 stimulation while decreasing pDC production of IFN-λ (**Fig. 4E**). The knock-down of *RAB5A* also reduced pDC production of IFN-α/λ and TNF (**Fig. 4F**). In contrast, *RAB7* knock-down (**Fig. 4C**) promoted the differentiation of HSCs into cDC1s and tended to increase their production of TNF upon R848 stimulation (**Fig. 4G**), without altering the development and activation of pDCs. *SEPTIN3* knock-down significantly increased the cDC1 output, and conversely reduced the pDC output; it also decreased pDC production of IFN-α/λ (**Fig. 4H**). To our knowledge, this is the first demonstration of the involvement of small GTPases in the development and activation of human cDC1s or pDCs.

**Fig. 4.**
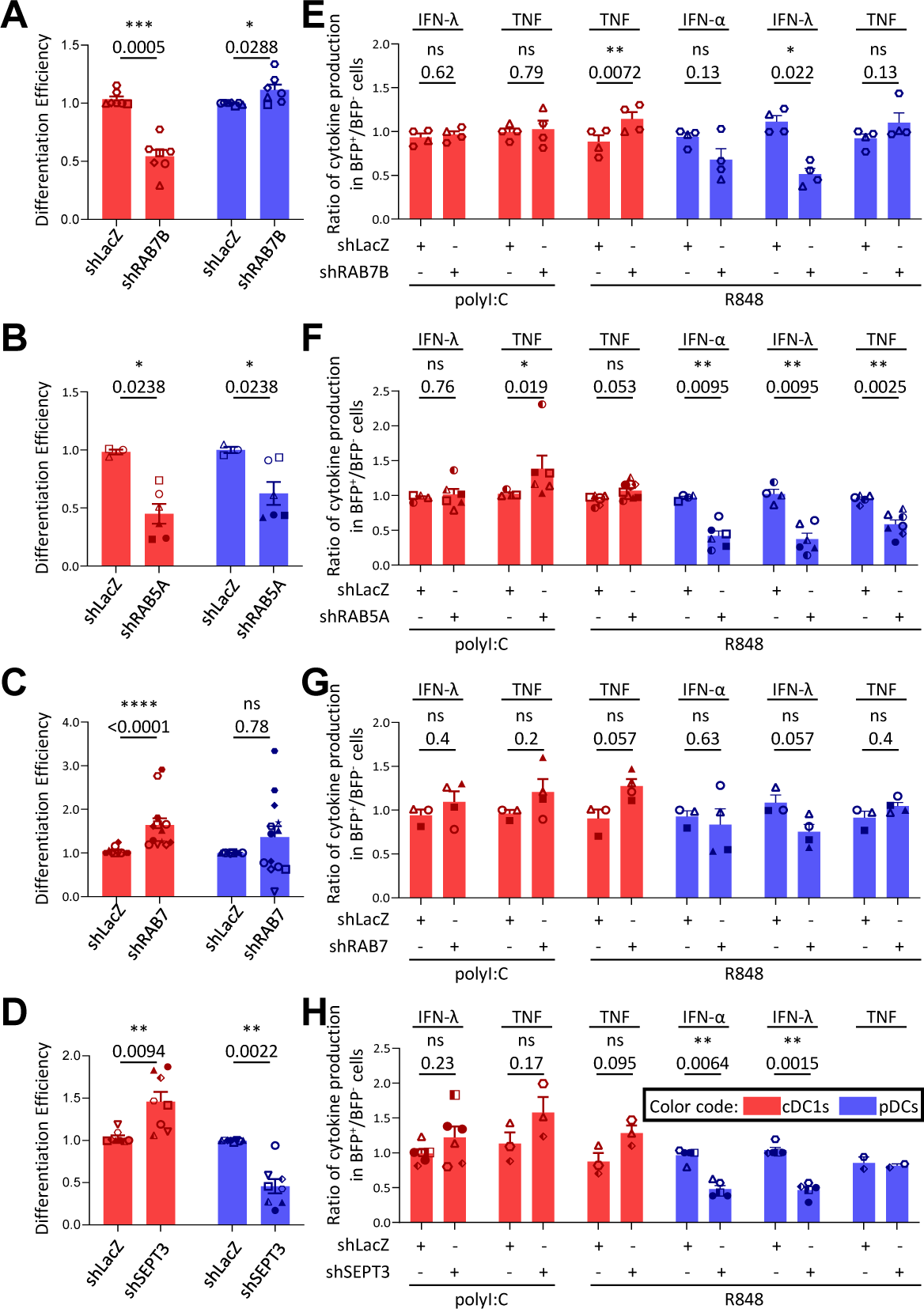
Identification of small GTPases controlling the development or activation of human cDC1s or pDCs. Impact on the development (**A-D**) of human cDC1 or pDC, and on their cytokine responses to TLR ligand stimulation (**E-H**), of the knock down of RAB7B (**A**, **E**), RAB5A (**B**, **F**), RAB7 (**C**, **G**) and SEPT3 (**D**, **H**). For the impact of RAB7B on DC development, seven independent experiments are shown, with four CB donors, and one shRNA; for RAB5A, three independent experiments, with two CB donors, and two different shRNA; for RAB7, ten independent experiments, with seven CB donors, and two different shRNA; for SEPT3, eight independent experiments, with six CB donors, and three different shRNA. For the impact of RAB7B on TLR response, four independent experiments are shown, with two CB donors, and one shRNA; for RAB5A, six independent experiments, with three CB donors, and three different shRNA; for RAB7, three independent experiments, with three CB donors, and two different shRNA; for SEPT3, six independent experiments, with four CB donors, and three different shRNA. For panel **A**, **D**, **E** and **H** statistical analyses were performed using a paired two-tailed t test. For panels **B**, **C**, **F** and **G**, statistical analyses were performed using a two-tailed non-parametric Mann-Whitney test. *, p<0.05; **, p<0.01; ***, p<0.001; ****, p<0.0001; ns, not significant.

### Identification of HIV-1 restriction factors in cDC1s/pDCs

MoDCs^58^ and cDC2s^11^ are resistant to infection by HIV-1 in a large part due to the restriction factor SAMHD1^59, 60^ which deprives the dNTP pool that is required for viral reverse transcription^61, 62^. HIV-1 restriction by SAMHD1 in MoDCs and cDC2s can be overcome by supplementing virus particles with Vpx from HIV-2/SIV^11, 58^, which targets SAMHD1 to proteasome for degradation^59, 60^. In contrast, supplementation by Vpx is insufficient to increase the susceptibility to HIV-1 infection of cDC1s and pDCs^11^, suggesting that these cells express other restriction factors. In cDC1s, RAB15 inhibits one of the earliest phases of the viral life cycle: the fusion of the viral envelope with the cell membrane^11^. RAB15 is also strongly expressed in pDCs, but its function in these cells remains unknown. Another restriction factor protecting cDC1s from HIV-1 infection may be palmitoyl-protein thioesterase 1 (PPT1), since it restricts infection of mouse cDC1s by vesicular stomatitis virus^57^. Thus, we examined whether SAMHD1, RAB15 and PPT1 contributed to the high resistance of cDC1s or pDCs to HIV-1 infection, also testing in parallel some of the genes studied in the previous sections. However, we first had to examine whether SAMHD1, RAB15 or PPT1 modulated the development or activation of human cDC1s or pDCs.

Unexpectedly, *SAMHD1* knock-down led to a strong and significant decrease in the cDC1 output of the cultures, whereas the pDC ouput was increased (**Fig. 5A**). To determine whether this effect could be due to loss of the dNTPase activity of SAMHD1, we tested whether similar results could be obtained by another method also leading to an increase in the cellular dNTP pool: the activation of their synthesis via the salvage pathway upon supplementation of the cultures with dNs. This did not seem to be affected the differentiation of cDC1s and pDCs as trongly as *SAMHD1* KD (**Fig. S4A**). Another mechanism through which *SAMHD1* knock-down could increase the pDC output of the cultures is via induction of spontaneous IFN-I responses^75^. Indeed, addition of exogenous IFN-I to human HSC cultures promotes their differentiation into pDCs^76^. This did not seem to be the case since pharmacological blockade of the receptor for IFN-I did not reverse the impact of *SAMHD1* knock-down on the cDC1 and pDC output of the cultures (**Fig. S4B**). Hence, how SAMHD1 controls the differentiation of human HSCs into cDC1s and pDCs remains to be deciphered. *SAMHD1* knock-down tended to increase cDC1 cytokine production, reaching significance for TNF in response to R848, whereas pDC cytokine production was unchanged, or decreased in the case of IFN-λ in response to R848 (**Fig. 5B**). *SAMHD1* knock-down slightly increased cDC1 susceptibility to HIV-1 infection, whereas it did not alter pDC permissiveness to the virus (**Fig. 5C**), consistent with the limited impact of the supplementation of HIV-1 particles with Vpx on their ability to infect cDC1s and pDCs^11^.

**Fig. 5.**
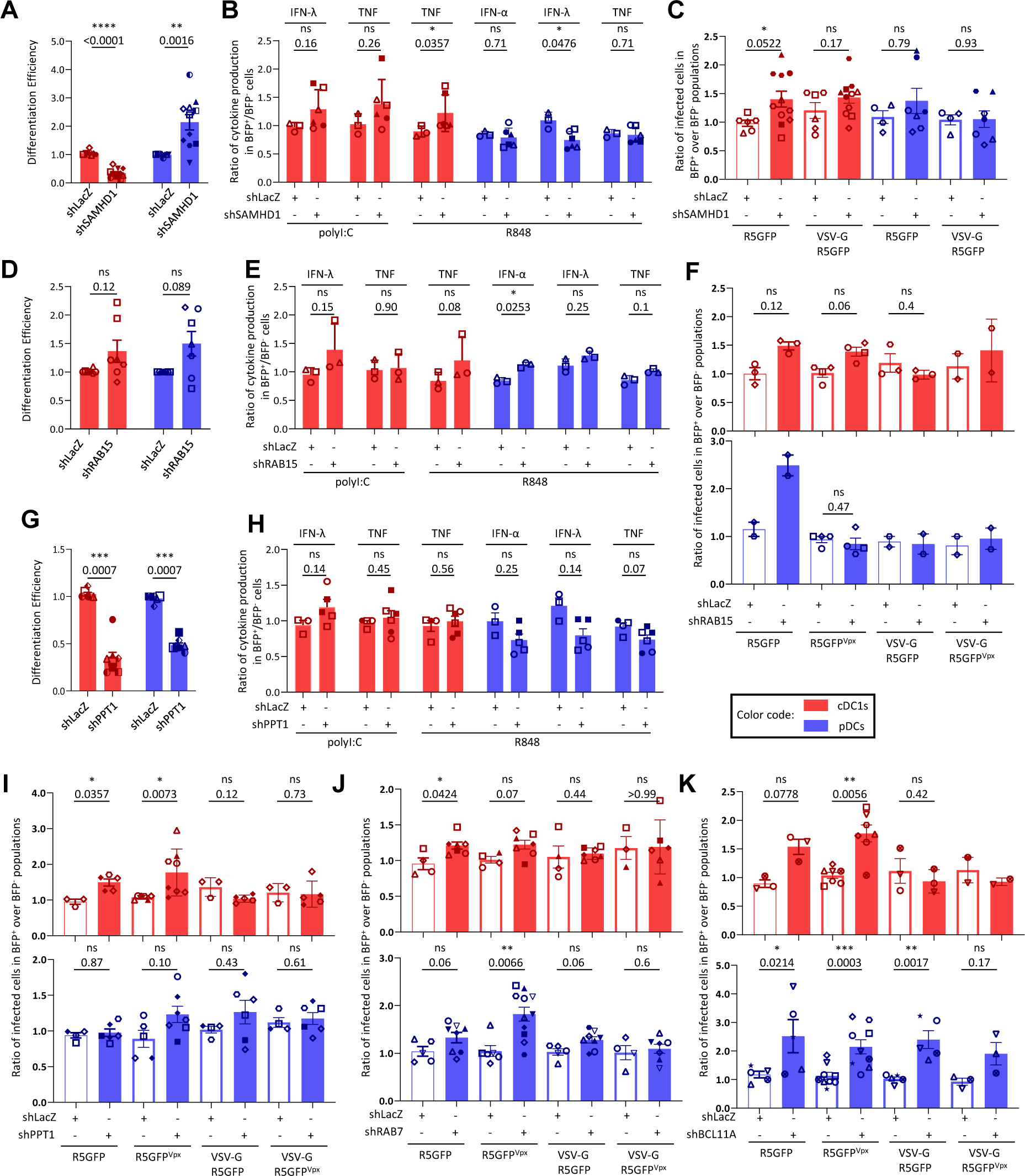
Identification of restriction factors controlling the development, activation or sensitivity to HIV-1 infection of human cDC1s or pDCs. Impact on cDC1 or pDC development (**A, D, G**), cytokine responses to TLR ligand stimulation (**B, E, H)**, and susceptibility to HIV-1 infection **(C, F, I, J, K)** of the knock-down of *SAMHD1* (**A-C**), *RAB15* (**D-F**), *PPT1* (**G-I**), *RAB7* (**J**), and *BCL11A* (**K**). For susceptibility to HIV-1 infection **(C, F, I, J, K),** HSCs were transduced with specific versus control (ShLacZ) shRNA lentivectors and differentiated for 2-3 weeks as indicated in Fig. 1. Differentiated cultures were then exposed to GFP-expressing HIV-1 viral particles (R5GFP), bearing different envelopes, with or without Vpx supplementation. After 48 hours of culture, the ratio of the percentages of infected (GFP^+^) cells within transduced (BFP^+^) versus non-transduced (BFP^-^) cDC1s or pDCs was measured. The results on DC development are shown from seven independent experiments, with five CB donors and one shRNA for *RAB15*; from eight independent experiments, with six CB donors and three different shRNA for *SAMHD1*; from six independent experiments, with four CB donors and three different shRNA for PPT1. The results on DC responses to TLR triggering are shown from three independent experiments, with three CB donors and one shRNA for *RAB15*; from three independent experiments, with three CB donors and two different shRNA for *SAMHD1*; from three independent experiments, with two CB donors and two different shRNA for *PPT1*. The results on susceptibility to HIV-1 infection are shown from four independent experiments, with four CB donors and one shRNA for *RAB15*; from six independent experiments, with six CB donors and two different shRNA for *RAB7*; from five independent experiments, with five CB donors and two different shRNA for *SAMHD1*; from six independent experiments, with five CB blood donors and two different shRNA for *PPT1*; from eight independent experiments, with eight CB donors and one shRNA for *BCL11A*. For panels **A**, **B**, **C**, **G**, **H**, **I** and **J**, statistical analyses were performed using a two-tailed non-parametric Mann-Whitney test. For panel **D**, **E**, **F**, and **K**, statistical analyses were performed using a paired two-tailed t test. *, p<0.05; **, p<0.01; ***, p<0.001; ****, p<0.0001; ns, not significant.

*RAB15* knock-down affected neither the development (**Fig. 5D**) nor the activation of human DCs except, for a slight increase in IFN-α production by pDCs (**Fig. 5E**). It tended to enhance cDC1 susceptibility to HIV-1 infection, regardless of the presence of Vpx (**Fig. 5F**), consistent with our previous observations^11^. *RAB15* knock-down increased pDC infection by HIV-1 R5GFP (**Fig. 5F**). However, unexpectedly, this increase was abolished by the presence of Vpx, implying a negative cross talk between RAB15 and SAMHD1 activities in pDCs. In both cDC1s and pDCs, the increase in HIV-1 infection induced by *RAB15* knock-down was abolished upon VSV-G-pseudo-typing of viral particles, showing that the antiviral function of RAB15 was linked to the endocytic route followed by the virus.

*PPT1* knock-down reduced the output of differentiated myeloid cells in our cultures, including cDC1s and pDCs (**Fig. 5G**), without appearing to impact overall cell expansion. *PPT1* knock-down did not affect the activation of cDC1s and pDCs (**Fig. 5H**). *PPT1* knock-down significantly increased the infection by HIV-1 R5GFP of cDC1s regardless of the presence of Vpx, but not of pDCs **(Fig. 5I**). This increase was abolished upon VSV-G-pseudo-typing of viral particles, showing that the antiviral function of PPT1 in cDC1s was linked to the endocytic route followed by the virus.

Among the other small GTPases that we studied, RAB7 tended to restrict HIV-1 infection in cDC1s and pDCs (**Fig. 5J)**, in a manner dependent of the virus endocytic entry route since the effects were reduced upon VSV-G-pseudo-typing of viral particles.

Finally, we observed that *BCL11A* knock-down enhanced HIV-1 infection of cDC1sand pDCs (**Fig. 5K**), irrespective of the presence of Vpx. Whereas this increase was abrogated in cDC1s upon VSV-G-pseudo-typing of viral particles, this did not appear to be the case in pDCs. The mechanisms underpinning the HIV-1 restriction activity of BCL11A in cDC1s may be due in part to its inhibition of CD4 expression on these cells (**Fig. S4C-D**).

## Discussion

Here we report a robust and relatively simple workflow to obtain large numbers of gene-silenced human cDC1s and pDCs. By utilizing the pipeline, for the first time to our knowledge, we extended to human cDC1s/pDCs the critical role of the transcription factors ID2 and DC-SCRIPT versus BCL11A in promoting the development of cDC1s versus pDCs as previously reported in mutant mice. Unexpectedly, we observed that *ID2* knock-down impaired pDC production of IFN-α/λ. The underlying mechanisms remain to be investigated.

Three small GTPases of the Rab family, RAB5A, RAB7 and RAB15, are expressed at similar levels between cDC1s and pDCs, but only RAB5A was required for their differentiation. The small GTPases RAB7B and SEPTIN3 are specifically expressed to high levels in cDC1s in both mice and humans, but only the former was required for the optimal development of human cDC1s in vitro from HSCs. The mechanisms by which small GTPases affect the development of cDC1s or pDCs remain to be investigated but may involve altered response to cytokines or growth factors due to deregulation of their receptor expression or signaling, since GTPase control intracellular trafficking including endocytosis/phagocytosis. Unexpectedly, *SEPTIN3* knock-down impaired pDC development and their production of IFN-α/λ, despite very low to undetectable expression of the gene in these cells. This suggested that SEPTIN3 might be transiently expressed in a proximal precursor of pDCs in a manner affecting their differentiation and the function of their progeny. More generally, pDC production of IFN-α/λ production seemed more frequently and profoundly impacted by the knock-down of the candidate genes tested than cDC1 cytokine production, suggesting more stringent requirements of various small GTPases for the building in pDCs of the specialized endosomes dedicated to signaling from TLR7/9 triggering to IFN-α/λ production.

Our pipeline not only allowed studying cDC1/pDC ontogeny and cytokine responses in response to TLR triggering; it was also suitable for screening candidate HIV-1 restriction factors and their mechanisms of action in human cDC1s and pDCs that are highly resistant to viral infections^11^. Here, we showed that BCL11A inhibits HIV-1 infection of pDCs and cDC1s, but probably through distinct mechanisms between these two cell types. Indeed, BCL11A-mediated inhibition of HIV-1 infection was dependent on the route of viral entry specifically in cDC1s, since it was abrogated upon VSVG-pseudo-typing of the virus, and *BCL11A* knock-down increased specifically on cDC1s the expression of CD4 which is the major entry receptor for HIV-1. Whether the upregulation of CD4 on cDC1s is directly induced by the transcription factor activity of BCL11A remains to be investigated. In contrast, the increase in the infection rate of pDCs upon *BCL11A* knock-down was not affected by the viral entry route or the presence of Vpx, suggesting an effect at later stages of the viral life cycle. BCL11B, a paralog of BCL11A, can suppress the initial phase of HIV-1 gene transcription in human microglial cells^77^ or of HIV-1 long terminal repeats in T cells^78^. Whether BCL11A can regulate the transcription of HIV-1 proviruses in pDCs remains to be investigated. Using our pipeline will also help understand how other viruses representing a threat for human health interact with human DC types.

Our work focused on using shRNA for target gene knock-down. Further applications of our pipeline could be achieving gene knock-down in primary human cDC1s/pDCs, by combining lentivector-based delivery of guide RNA with electroporation-based delivery of Cas protein, as was previously achieved in other experimental systems of in vitro differentiation of human immune cells from HSCs^79, 80^.

In conclusion, we were successful in setting-up a robust pipeline to genetically edit human cDC1s and pDCs, enabling to test the impact of the knock-down of candidate genes on their biology. This allowed us identifying novel mechanisms controlling human cDC1/pDC development, cytokine production or susceptibility to HIV-1 infection. This technological breakthrough should allow conducting larger scale genetic screens to advance our knowledge on the functions of human cDC1s/pDCs and their molecular regulation, with the perspective to manipulate them for designing complementary approaches to boost existing vaccination or immunotherapies against viral infections or cancer.

## Acknowledgements

We thank the staff of the CIML flow cytometry core facility. This work was supported by CNRS, INSERM, Agence Nationale de Recherches sur le SIDA et les Hépatites Virales (ANRS, project ECTZ25472, to M.D.). X.Luo received fellowships from SIDACTION and ANRS.

## Author Contributions Statement

X.Luo. designed, performed and analysed the experiments, wrote the manuscript. A.J. contributed help to some of the experiments. X. Lahaye and P.M. contributed advice and key reagents. N.M. contributed to direct and fund the study, and gave advice on experiments. M.D. directed and funded the study, designed and help analysing experiments, and wrote the manuscript. All authors contributed to edit the manuscript.

## Methods

### Cells

Human CB CD34^+^ HSCs were bought from ABCell-bio, France. For each purchasing campaign, a tube of 200,000 cells was first ordered for a series of donors, to test their differentiation efficacy into cDC1s and pDCs, before ordering additional tubes of between 300,000 and 700,000 cells for the donors yielding the most efficient differentiation. HEK293FT cells were cultured in DMEM medium, GlutaMAX (Thermo Fisher 61965-026) complemented with FBS 10% (Thermo Fisher 10270-106) and Penicillin/Streptomycin. THP-1 cells (ATCC) were cultured in RPMI 1640 medium, GlutaMAX (Thermo Fisher 61870-010) complemented with FBS 10% (Thermo Fisher 10270-106) and Penicillin/Streptomycin. OP9 and OP9-DL1 cells were cultured as previously described^31, 81^.

### Lentivector production and concentration

pLKO.1-GFP plasmid was adapted to pLKO.1-mTagBFP by replacing the CDS of GFP with that of mTagBFP. Briefly, CDS of mTagBFP was cloned from pTRIF-SFFV-mTagBFP-2A plasmid^82^ by using the following primers: mTagBFP-F: cagggggatccaccggagcttaccATGAGCGAGCTGATTAAGGAG and mTagBFP-R: taccgatgcatggggtcgtgcgctcctttcggtcgggcgctgcgggtcgtgg-ggcgggcgTTAATTAAGCTTGTGCCCCAGT. The amplified fragment and pLKO.1-GFP plasmid were digested with NsiI and BamHI at 37℃ overnight. Digested fragment and plasmid were then ligated and used to transform competent E. coli cells. Positive clones were selected and verified by using bacterial-liquid PCR and sequencing. mTagBFP expression measured by flow cytometry in HEK293FT cells transfected with the plasmid. The packaging ability of the lentivectors was also verified to ensure that it did not differ from that of the parental pLKO.1-GFP vectors. HEK293FT cells were seeded into 6-well plates. The transfection was performed by combining 0.3μg pVSV-G, 0.9μg pPAX2 (psPAX2 was a gift from Didier Trono (Addgene plasmid # 12260; http://n2t.net/addgene:12260; RRID:Addgene_12260)) and 1.05μg LKO.1-shRNA-BFP as previously described^11^. The target sequences for the shRNA that were selected to be used in experiments are listed in **Table S1**. The next day, the supernatant was discarded and replaced by 3ml of X-VIVO 15 serum-free medium. Two days after, the supernatant was harvested into 50ml Falcon tubes. A small aliquot of the virus (500µl) was used for titration. The bulk of the supernatant was stored at −80°C until the lentivector titration was known, then it was thawed for concentration via ultracentrifugation after the lentivectors were titrated. To construct the plasmid for IRF8 overexpression, CDS of IRF8 from IRF8 human tagged ORF clone (RG217646, Origene) were cloned and digested with BamHD1 and XhoI. The fragment was then cloned into TRIP-SFFV-mTagBFP-2A (Addgene, #102585). To produce lentivectors for IRF8 overexpression, transfection of previously seeded HEK293FT cells was performed by combining 0.3μg pVSV-G, 0.9μg psPAX2 and 1.05μg pTRIF-SFFV-IRF8-mTagBFP or empty vector for negative control. For titration, briefly, serial dilutions of the lentivectors were made and added to previously prepared HEK293FT cells plated in 96 well flat bottom plates at 5,000 cells/well. Forty-eight hours after, cells were harvested by Trypsin digestion and resuspended into PBS. The infection rate in each well were measured by FACS. The titers of the lentivectors were calculated as previously reported^83, 84^, as infectious units (IU) per mL, based on the percentages of BFP^+^ or GFP^+^ cells and on Poisson distribution (-ln(100-% BFP^+^ or GFP^+^)/100); only values ranging between 1% and 40% were used for calculation to ensure that they felled into a linear distribution. For concentrating them, the lentivectors were thawed and ultracentrifugated at 19,000rpm for 2 hours at 4°C. The supernatant was then discarded and the pellets resuspended with the volume calculated to achieve a final lentivector titer of 10^7^ IU/ml. The resuspended concentrated lentivectors were then aliquoted at 100μl/tube and stored at −80°C until use. The lists of the plasmids used/generated is given as **Table S2**.

**Table S1.**
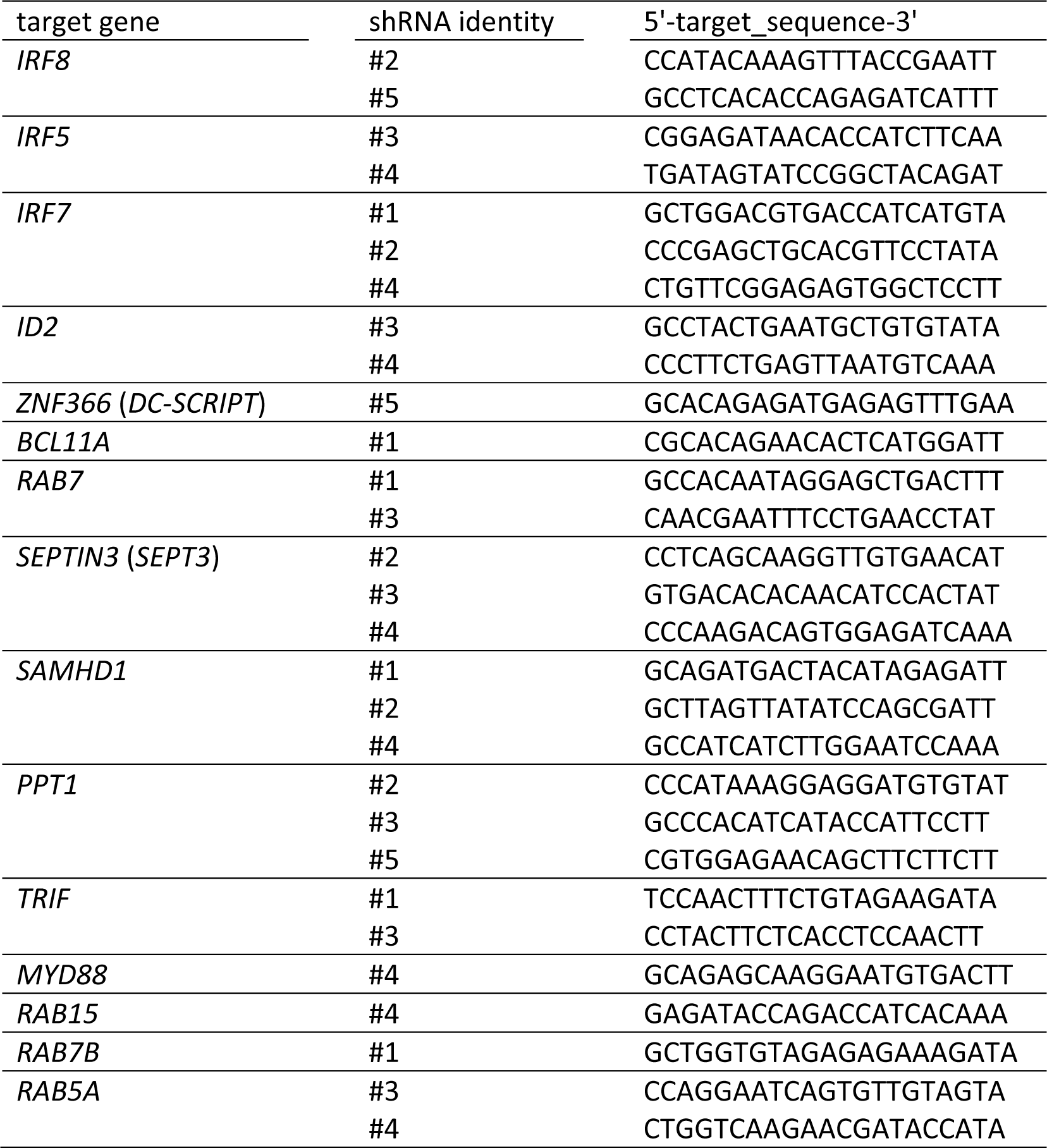
Target sequences for the shRNA selected for use in experiments.

**Table S2.**
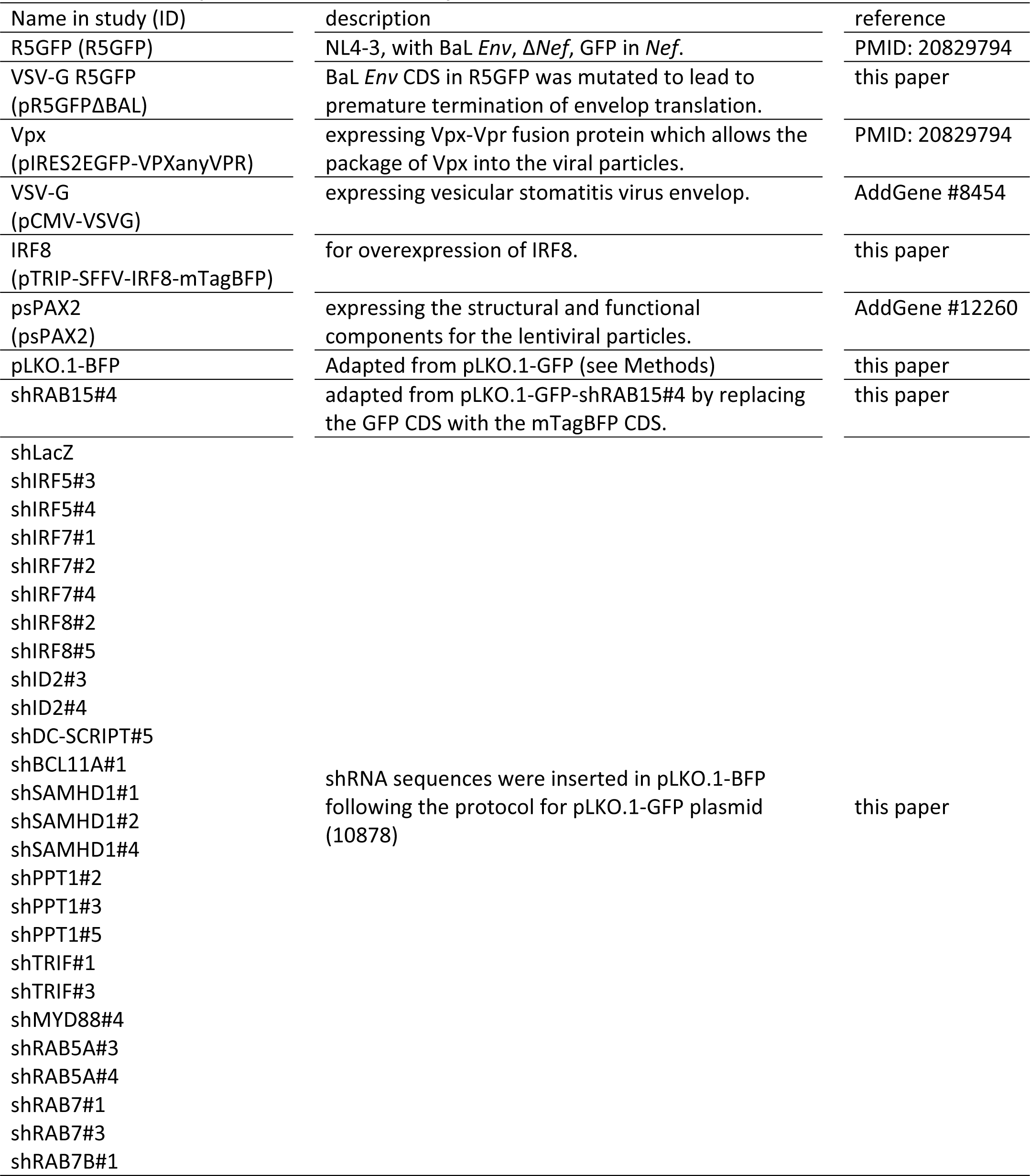
List of the plasmids used in the study.

### Transduction of THP-1 cells

Cultured THP-1 cells were harvested and resuspended into 10^6^ per ml. 500μl of the cells were seeded into 24 well plate and 500μl of nonconcentrated lentivectors were added, complemented with polybrene, the transduction was performed via spinoculation at 800g, 2 hours under room temperature. After spinoculation, cells were put back into the incubator and cultured for one week. The cells were finally cultured in 90mm petri dish before harvest due to the quick expansion.

### RNA extraction and qRT-PCR

The KD efficiency was assessed by qRT-PCR for the genes against which we did not identify commercially available antibodies of good specificity that worked in flow cytometry or western blot. The target genes tested and primer pairs used are listed in **Table S3**. Before RNA extraction, the transduction rate was measured by flow cytometry to ensure that enough percentage of cells are transduced (>60%). The cells were then pelleted into a RNase-free Eppendorf tubes and were lysed in 1ml Trizol reagent (Invitrogen, Ref. 15596026). 10 minutes after the addition of Trizol, 200μl chloroform were then added. The mixtures were then thoroughly mixed and keep them in room temperature before centrifugation at 12000 rpm at 4°C for 15 mins. The upper clear fractions were then transferred to new RNase free eppendorf tubes (around 450μl) and 500μl isopropanol were then added and thoroughly mixed. The tubes were kept at room temperature for another 10 minutes before centrifugation at 12000rmp under 4℃ for 10 mins. After the centrifugation, the supernatant was discarded and the pellets were washed with 1ml 70% ethanol by centrifugation at 12000rmp at 4°C for 5 mins. The supernatant was then discarded and the pellets were then dried under an RNase-free chemical hood for 20 mins and 100μl RNase-free water were not added until the residual liquid could not be observed. After complete dissolve, the concentration was measured by NanoDrop 2000/2000c Spectrophotometer (Thermo fisher) and the concentrations of RNA were finally adjusted to 100ng/ul. cDNA was prepared from 2μg total RNA from each sample under the instruction of QuantiTech reverse transcription Kit (Qiagen, 205311). Samples for QPCR were then prepared under the instruction of ONEGreen fast qPCR kit (Ozyme, OZYA008-40) and loaded on ABI 7500 fast Real-time PCR system. The relative mRNA of each gene was normalized to the mRNA level of GAPDH in each sample.

**Table S3.**
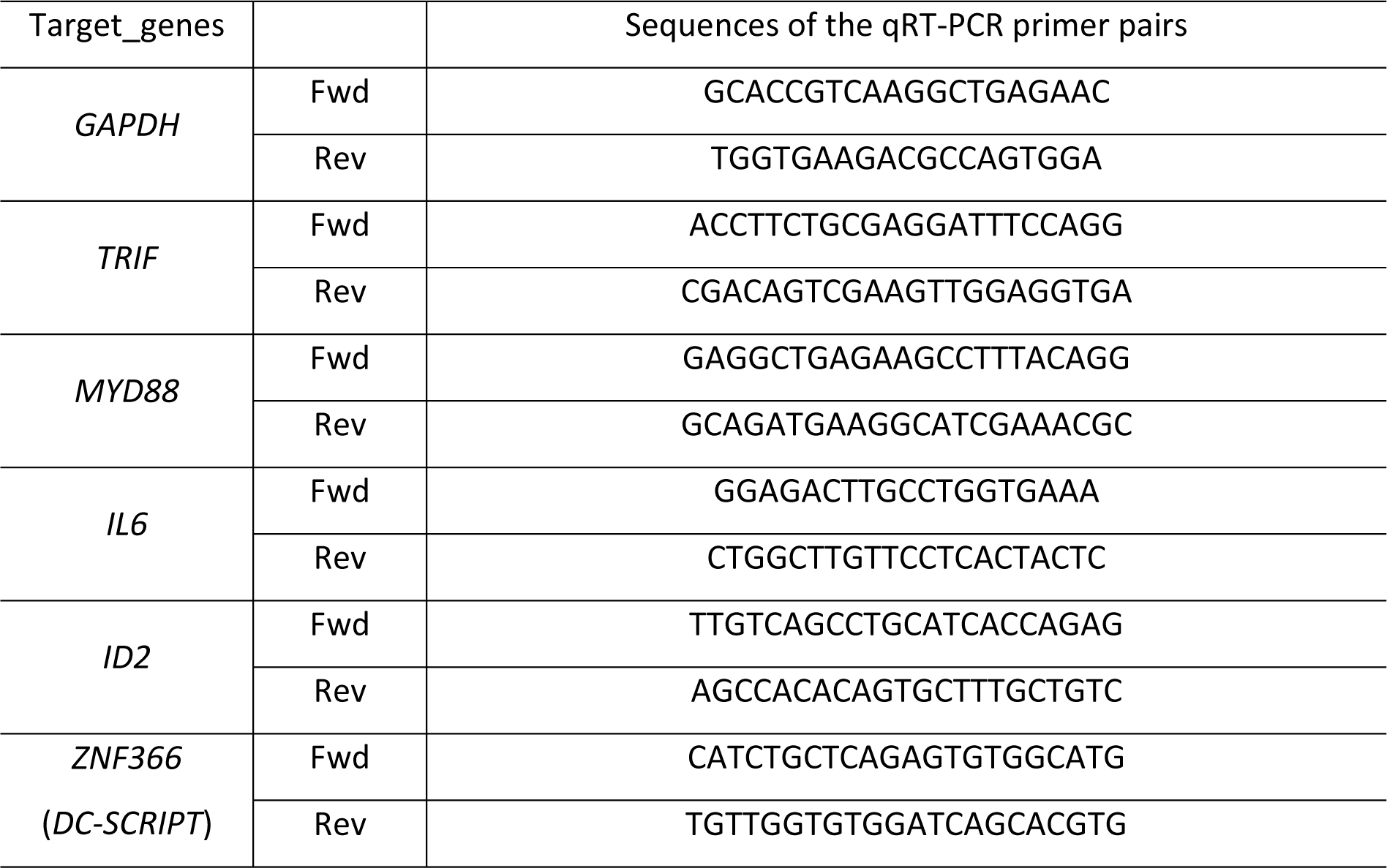
Primers used for qRT-PCR.

### Western blot

For analysis the knocking down efficiency of the shRNA candidates for different genes including SAMHD1, RAB7B, RAB7, RAB5A, PPT1, BCL11A, IRF8 and IRF5, shRNA transduced THP-1 cell lysates were prepared, while for SEPT3, shRNA transduced HEK293FT cell lysates were prepared. 5×10^6^ cells were pelleted after detection of the transduction rate by flow cytometry and were washed in PBS and lysed in 200µl of Pierce RIPA buffer (Thermo Scientific, 89900), complemented with protease inhibitor (Roche; 1187358001) on ice for 30 mins. 40µl of 6X Laemmli buffer (4% SDS, 20% Glycerol, 0.125M Tris-HCl pH6.8, 10% 2-mercaptoehtanol, 1% bromophenol blue) was added and samples were boiled at 95°C for 20min. 40μl of cell lysates were loaded on previously-prepared 10% SDS-PAGE gels and transferred on nitrocellulose membrane after the membrane was saturated 30min at room temperature with transferring buffer. The membranes were then blocked with 0.5% (w/v) non-fat milk powder in TBS buffer. Proteins were blotted with antibodies against SAMHD1 (1:1000), RAB7B (1:1000), RAB7 (1:1000), RAB5A (1:1000), PPT1 (1:1000), BCL11A (1:1000), IRF8 (1:1000) and IRF5 (1:1000) and actin (1:10000) (**Table S4**) in 0.25% (w/v) non-fat milk powder in TBS overnight at 4°C in TBS buffer. Membranes were washed three times 10min in TBS with 0.1% Tween (TBS-T) and incubated for 1hr with secondary goat antibodies against mouse (1/10000) or donkey antibodies against rabbit (1/5000) in 0.25% (w/v) non-fat milk powder in TBS buffer. ECL signal was obtained with Pierce western blot substrate (Thermo Scientific, 32106) and was recorded on the ChemiDoc XRS Imager (Biorad).

**Table S4.** Antibodies used for western blot.

| Antigen | dilutions | species | brand | reference | RRID |
| --- | --- | --- | --- | --- | --- |
| RAB7 | 1:1000 | Rabbit IgG | Proteintech | 55469-1-AP | AB_11182831 |
| RAB7B | 1:1000 | Mouse | Fisher Scientific | 16140114 | Not available |
| RAB5A | 1:1000 | Rabbit IgG | Proteintech | 11947-1-AP | AB_2269388 |
| SEPT3 | 1:1000 | Rabbit IgG | Thermofisher | PA5-31124 | AB_2548598 |
| PPT1 | 1:1000 | Rabbit | Sigma aldrich | HPA021546 | AB_1855667 |
| IRF5 | 1:1000 | Rabbit IgG | Proteintech | 10547-1-AP | AB_2280474 |
| IRF8 | 1:1000 | Rabbit IgG | Proteintech | 18977-1-AP | AB_10596482 |
| BCL11A | 1:1000 | Rabbit IgG | Thermofisher | MA5-34678 | AB_2848586 |
| SAMHD1 | 1:1000 | Rabbit | C.S.T | 49158S | Not available |
| Actin | 1:10000 | mouse | C.S.T | 3700S | Not available |

### Expansion and transduction of HSCs followed by their differentiation into cDC1s and pDCs

HSCs were amplified and differentiated into cDC1s and pDCs as reported previously^31, 81^, with the specific adaptations described here in the result section and shown in **Extended Data Fig. 1**. The transduction protocol was divided into three steps. 1) The first step was the “priming phase”. One vial of expanded HSCs (100,000-400,000 cells) was thawed and suspended into 1ml X-VIVO 15 serum free medium and put into one well of a 24-well plate. Cytokines (FST, meaning FLT3-L, SCF and TPO) were added at the final concentration of 100 ng/ml. Cells were then incubated at 37°C, 5% CO2 for 24 hours. 2) The second step was the transduction itself. The pre-stimulated cells were harvested and resuspended in X-VIVO 15 serum-free medium at the concentration of 1 to 2×10^5^/100μl. 100μl of the cell suspension was seeded per well of a 96-well U-bottom plate. Then, each well received 100μl of concentrated lentivectors (∼10^7^ IU/ml in X-VIVO 15 serum free medium supplemented with FST to reach the same final concentration as for the priming phase). Finally, poxamer 407 (Sigma) was added to each well, at a final concentration of 100μg/ml. 3) The third step consisted in seeding the cells to induced their differentiation into cDC1s/pDCs. 24 hours after step 2, the transduced cells were harvested and counted. 2×10^4^ cells were seeded onto the previously prepared OP9/OP9-DL1 feeder layer, in 24 well plates, as previously reported^31, 81^. Half of the supernatant was changed once a week with fresh medium containing FST, as previously described^31, 81^. Two to three weeks after initiation of the differentiation phase, the cells were harvested for analysis of the proportion of cDC1s/pDCs or for performing TLR stimulation or HIV-1 infection experiments.

### Flow cytometry

Cells were harvested and filtered through a 70μm cell-strainer to obtain single cell suspensions. The cells were first blocked with human Fc block (BD Biosciences) and then stained with live dead fixable aqua dead kit (Invitrogen) and the antibodies listed in **Table S5** in staining buffer (PBS containing 2% FCS). The purified anti-IFN-λ antibody was conjugated in-house with Alexa Fluor 647 by following the instructions of the antibody labeling kit (Invitrogen, A20186). The obtained labeled anti-IFN-λ antibody solution was used at 1d200 for staining. Intracellular staining was performed using a Fix/Perm kit (BD Biosciences). Samples were acquired on LSR II or Fortessa (BD Biosciences) and analyzed with FlowJo (Tree Star) software.

**Table S5.**
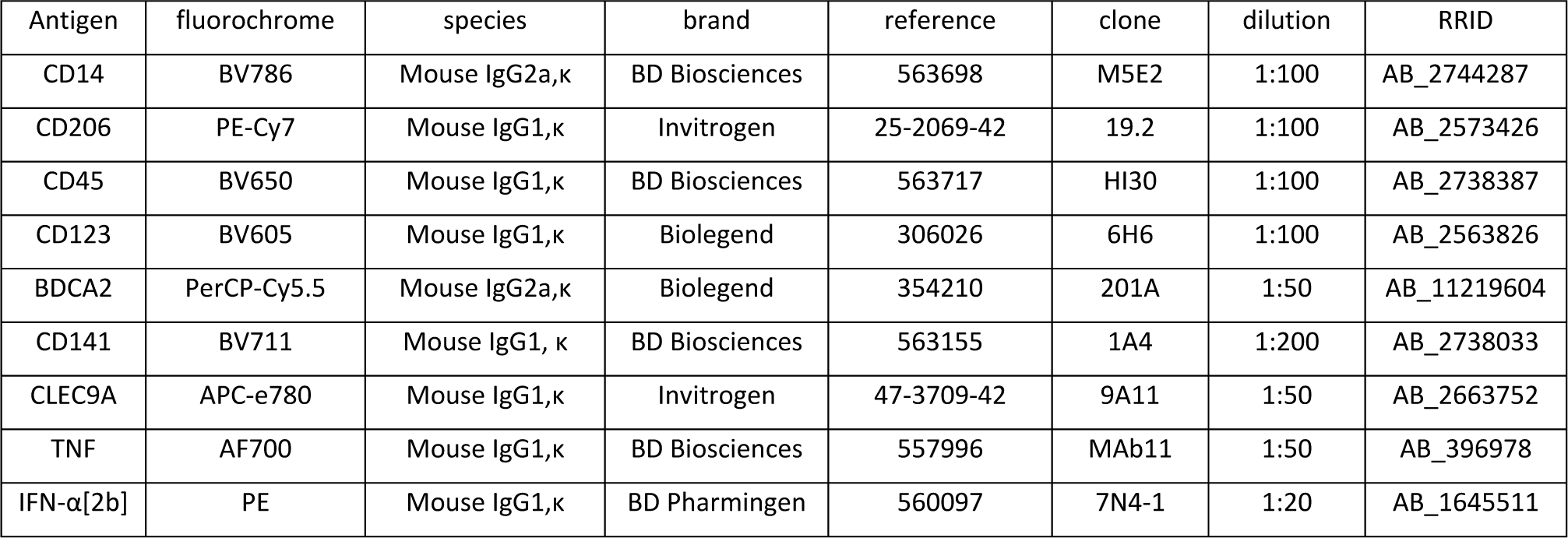
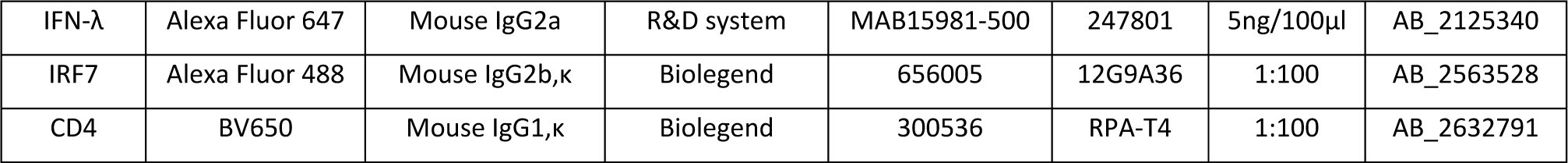
Antibodies used for flow cytometry

### Definition of “differentiation efficacy”

To define the “differentiation efficacy” of cDC1s and pDCs, we first calculated the ratio of the percentages of total BFP^+^ cells to the total BFP^-^ cells as “ratio_total” in all live and CD45^+^ cells. The ratio of the percentages of BFP^+^ pDCs to BFP^-^ pDCs as “ratio_pDCs” and that of cDC1s as “ratio_cDC1s”. The pDC differentiation efficacy was then calculated by dividing ratio_pDCs with ratio_total, and cDC1 differentiation efficacy was calculated by dividing ratio_cDC1 with ratio_total. In most of the experiments, values of differentiation efficacy from shLacZ-transduced samples were around 1 as expected for lack of specific effect of transduction in pDCs or cDC1s as compared to the other CD45^+^ cells in the cultures. However, due to the donor differences or fluctuations in the culture system, values of differentiation efficacy from shLacZ-transduced samples were sometimes as low as 0.5 or as high as 1.5. In these cases, these values were normalized to 1 and the values for the test shRNA-transduced samples in the same experiments were normalized consistently.

### Stimulation with synthetic TLR ligands

To activate DCs, 5μg/ml R848 or 10μg/ml polyI:C were added to the culture, followed one hour later by the addition of BFA to block exocytosis and hence enable intracellular detection of cytokines by flow cytometry. Five hours after the addition of the TLR ligands, cells were harvested and stained for FACS detection of TNF, IFN-α or IFN-λ in cDC1s and pDCs. The effect of gene knock-down on cDC1 or pDC activation was assessed by calculating the ratio of the percentage of cytokine-producing cells within BFP^+^ cDC1s/pDCs to that within BFP^-^ cDC1s/pDCs.

### Stimulation with synthetic TLR ligands

To activate DCs, 5μg/ml R848 or 10μg/ml polyI:C were added to the culture, followed one hour later by the addition of BFA to block exocytosis and hence enable intracellular detection of cytokines by flow cytometry. Five hours after the addition of the TLR ligands, cells were harvested and stained for FACS detection of TNF, IFN-α or IFN-λ in cDC1s and pDCs. The effect of gene knock-down on cDC1 or pDC activation was assessed by calculating the ratio of the percentage of cytokine-producing cells within BFP^+^ cDC1s/pDCs to the percentage of cytokine-producing cells within BFP^-^ cDC1s/pDCs.

### HIV-1 production and infection

All species of HIV-1 were produced by transfecting HEK293FT cells that were seeded in 6-well plates in BSL-3, using the TransIT-293 Transfection Reagent (Mirus Bio, USA) and following the instruction of the manufacturer. For the production of HIV-1 R5GFP, 2.25μg pR5GFP were used for each well. For HIV-1 R5GFP^Vpx^, 0.3μg pIRES2EGFP-VPXanyVpr+ 1.75μg pR5GFP were mixed for each well. To mutate the wild-type envelop (BAL) on pR5GFP, overlap PCR was used to introduce TA right after the N-terminal 249 bases in order to create a stop codon with the sequential G nucleoside and hence terminate the translation process. The pR5GFP with deficient envelop gene were named pR5GFP^ΔBAL^ and verified for their inability to produce infectious viral particles. For the production of VSV-G pseudo typed R5GFP^ΔBAL^, 0.3μg pCMV-VSV-G+ 1.95μg pR5GFP^ΔBAL^ were mixed for each well. For VSV-G pseudo typed R5GFP^ΔBAL+Vpx^, 0.3μg pCMV-VSV-G+0.3μg pIRES2EGFP-VPXanyVpr+1.65μg pR5GFP^ΔBAL^ were mixed for each well. The next day after the transfection, the supernatant were replaced with 3ml pre-warmed differentiation medium (α-MEM Glutamax, 10% FCS, 1% penicillin-streptomycin, 1X NEAA, 2mM L-Glutamine, 10mM HEPES, 1mM sodium pyruvate and 50μM β-mercaptoethanol). The virus containing supernatant were harvested at two days post-transfection, filtered at 45μm, aliquoted into 1ml Eppendorf tubes and stored at −80℃. For the infection of *in vitro* differentiated DC cultures in 24 well plates, 500μl of supernatant were removed and 320μl fresh medium supplemented with cytokine cocktails for the differentiation were added. 180μl of virus were then added, with protamine supplementation at a final concentration of 1μg/ml. Infection was performed under spinoculation at 800g for two hours. Plates were put back into the incubator for 48 hours before staining for assessing by flow cytometry the infection rate of the different cell types.

### Blockade of the IFN-I response

Vaccinia virus B18R carrier-free recombinant protein (eBioscience, 34-8185-81) was purchased and used at 3μg/ml as a type I IFN blocker, as previously described^11^.

### Bulk-RNA sequencing

The differentiated cDC1s and pDCs from two CB donors were sorted by flow cytometry to a purity>98%. The concentration and integrity of each RNA sample was verified by using Agilent RNA 6000 Nano chips and analyzing them with the 2100 bioanalyzer according to the manufacturer instructions. Bulk RNA sequencing was then performed by the sequencing core facility of the Curie Institute in Paris and the results analyzed as previously described^11^.

### Statistical analysis

No statistical methods were used to pre-determine sample sizes but our sample sizes are similar to those reported in previous publications^11, 31^. Data distribution was assumed to be normal but this was not formally tested. All quantifications were performed with awareness of experimental conditions, meaning not in a blinded fashion. No data were excluded. Statistical parameters including the definitions and exact value of n (number of biological replicates and total number of experiments), and the types of the statistical tests are reported in the figures and corresponding legends. Statistical analyses were performed using Prism v8.1.2 (GraphPad Software). Statistical analysis was conducted on data with at least three biological replicates for the test shRNA conditions. Comparisons between groups were planned before statistical testing and target effect sizes were not predetermined. Error bars displayed on graphs represent the mean±SEM. A two-tailed non-parametric Mann-Whitney test was used when the same shRNA control condition was used for assessing the effect of distinct shRNA targeting the same candidate gene, for each donor. A paired two-tailed t test was used when only one shRNA was used for the candidate gene, and hence the numbers of control and test conditions were identical and could be paired without repeating the same control values. Statistical significance was defined as * for p<0.05, ** for p<0.01, *** for p<0.001 and **** for p<0.0001.

### Data availability

RNA-seq data will be deposited in the Gene Expression Omnibus (GEO) repository once the manuscript has been accepted in principle. All other data generated or analysed during this study are included in this published article (and its supplementary information files).

## Supplementary information

**Supplementary Fig. 1.**
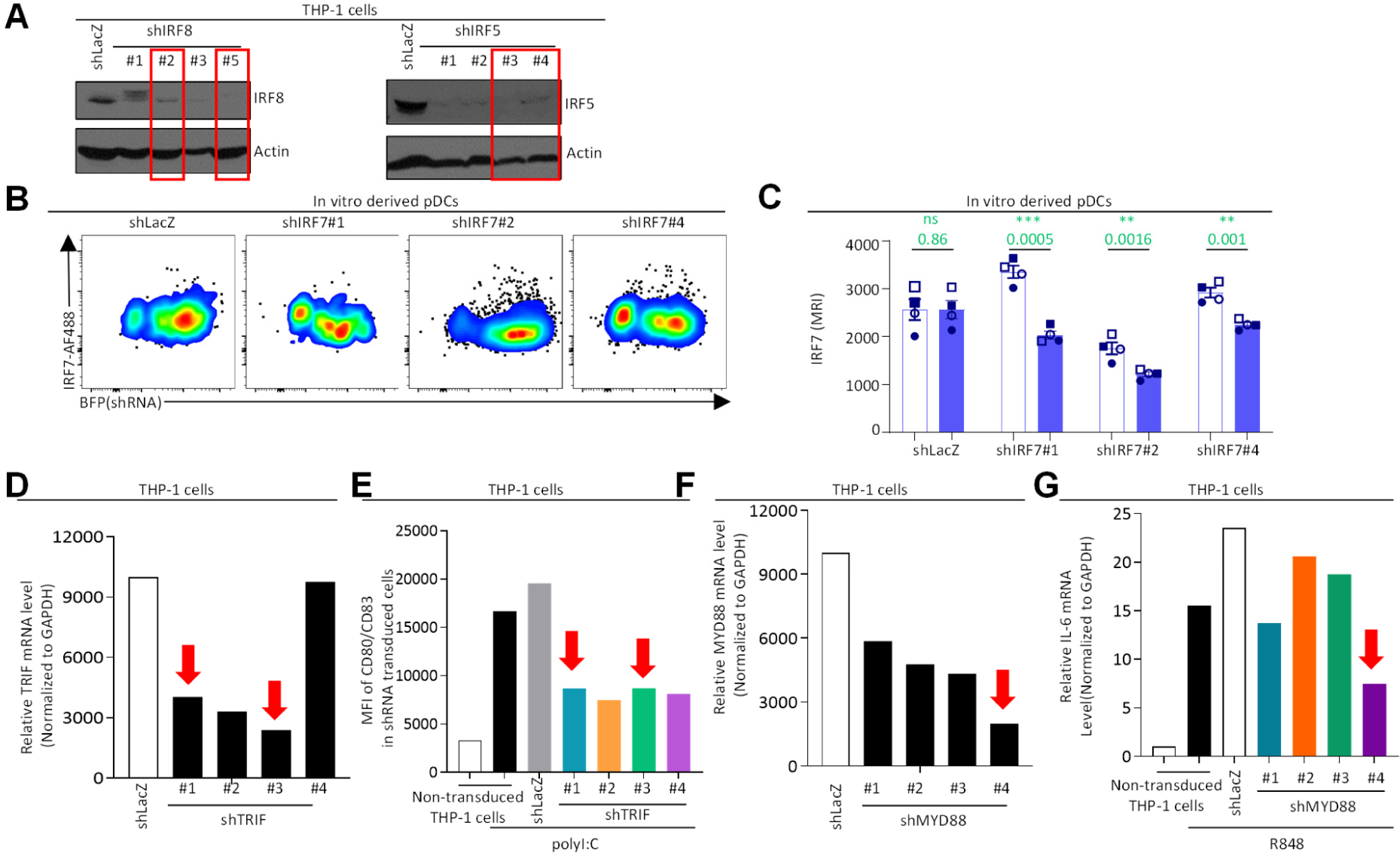
Evaluation of the knock-down efficiency for IRF8, IRF5, IRF7, TRIF and MYD88. (**A**) Knocking down efficiency with the lentivectors shIRF8 or shIRF5, compared with control (shLacZ), as assessed by Western blot on transduced THP-1 cells. (**B**) Results of one representative experiment assessing the knock-down efficiency with the lentivectors shIRF7 on pDCs by FACS. (**C**) IRF7 MFI of transduced and non-transduced pDCs, across four independent experiments. (**D-E**) *TRIF* knock-down efficiency was assessed in THP-1 cells at the mRNA level by qRT-PCR (**D**), and at the functional level by measuring by FACS CD80/CD83 upregulation upon polyI:C stimulation (**E**). (**F-G**) *MYD88* knock-down efficiency was assessed in THP-1 cells at the mRNA level by qRT-PCR (**F**), and at the functional level by measuring by qRT-PCR *IL6* mRNA upregulation upon R848 stimulation (**G**). In all instances, control (shLacZ) lentivectors were used for comparison. All shRNA tested against IRF7 where used in the experiments. For the other target genes, red arrows indicate the shRNA that were selected for functional assays in the DC cultures. For panel C, statistical analyses were performed using a two-tailed paired t test. *, p<0.05; **, p<0.01; ***, p<0.001; ****, p<0.0001; ns, not significant.

**Supplementary Fig. 2.**
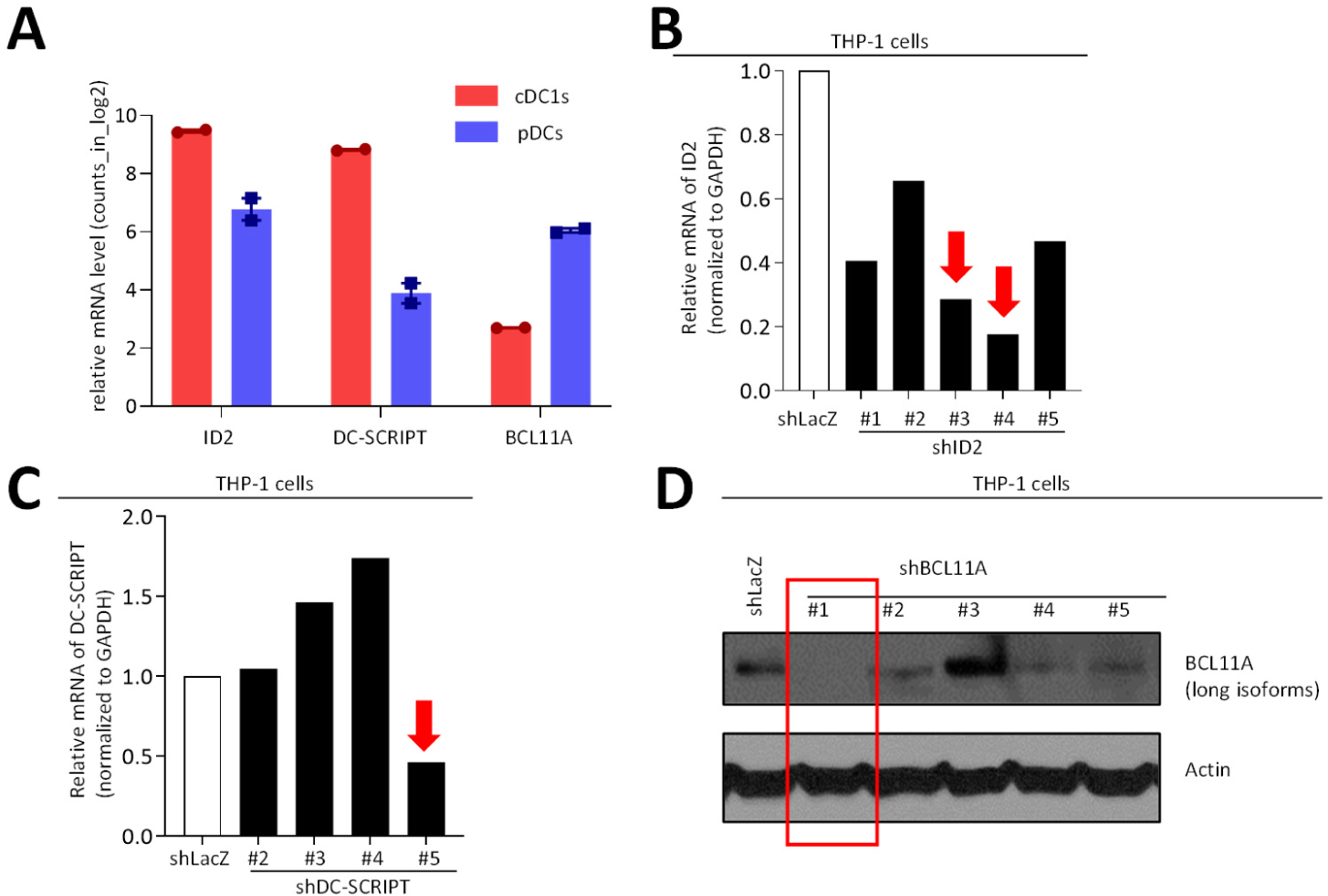
Evaluation of the knock-down efficiency for candidate transcription factors. (**A**) Relative mRNA level of candidate transcription factors ID2, DC-SCRIPT and BCL11A in cDC1s and pDCs. The data shown are from one RNA-seq bulk experiment performed with cells isolated from two different donors. (**B-C**) Knock-down efficiency was assessed on THP-1 cells by qRT-PCR for *ID2* (**B**) and *DC-SCRIPT* (**C**). (**D**) *BCL11A* knock-down efficiency was assessed in THP-1 cells at the protein level by western blot. (**B-D**) The results shown are from one experiment representative of at least two independent ones. Red arrows and boxes indicate the shRNA that were selected for functional assays in DC cultures, based on the strength and reproducibility of their knock-down efficiency.

**Supplementary Fig. 3.**
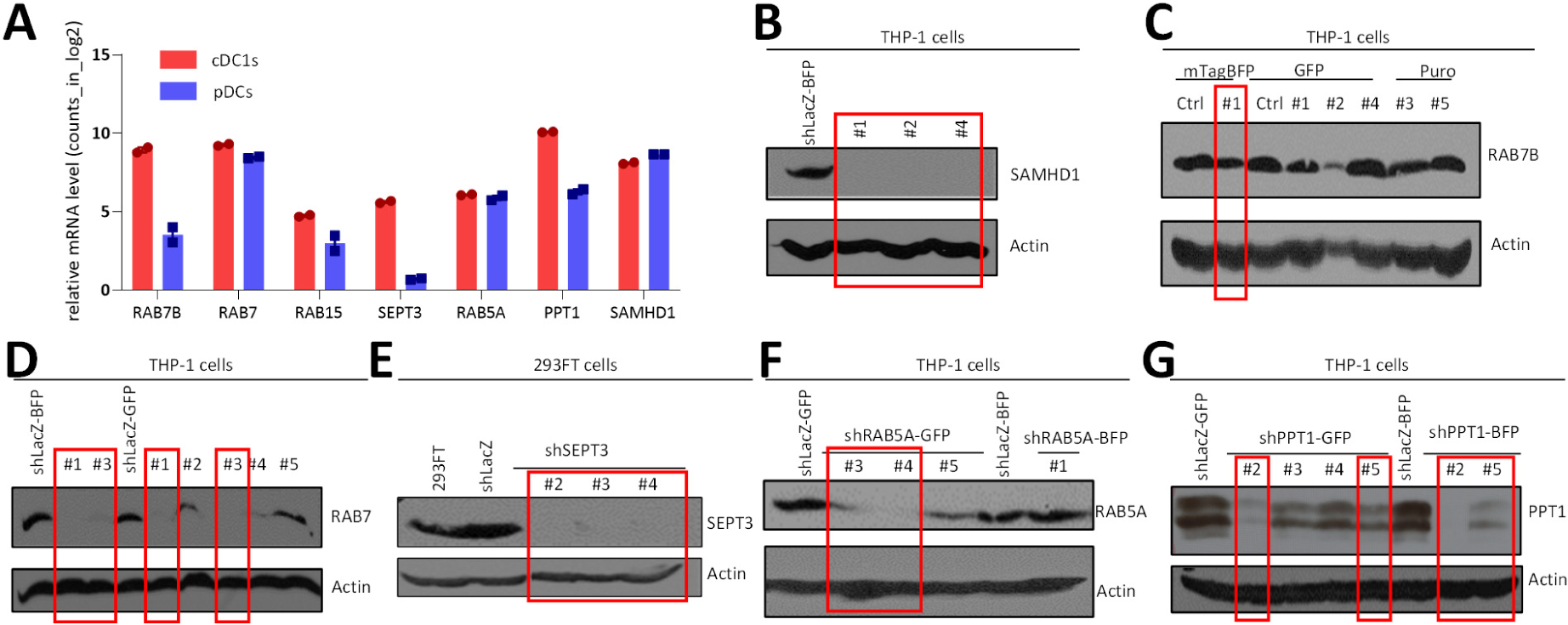
Evaluation of the knock-down efficiency of shRNAs for candidate small GTPases. **A**) Relative mRNA level of candidate genes in cDC1s and pDCs. The data shown are from one RNA-seq bulk experiment performed with cells isolated from two different donors. (**B-G**) Knock-down efficiency assessed at the protein level by western blot for SAMHD1 (**B**), RAB7B (**C**), RAB7 (**D**), SEPT3 (**E**), RAB5A (**F**) and PPT1 (**G**). The results shown for each target gene are from one experiment representative of at least two independent ones. Efficacy of KD was assessed in THP-1 cells except for SEPT3 expression of this protein was not detected in these cells such that HEK293FT cells were used instead. Red boxes indicate the shRNA that were selected for functional assays in DC cultures, based on the strength and reproducibility of their knock-down efficiency.

**Supplementary Fig. 4.**
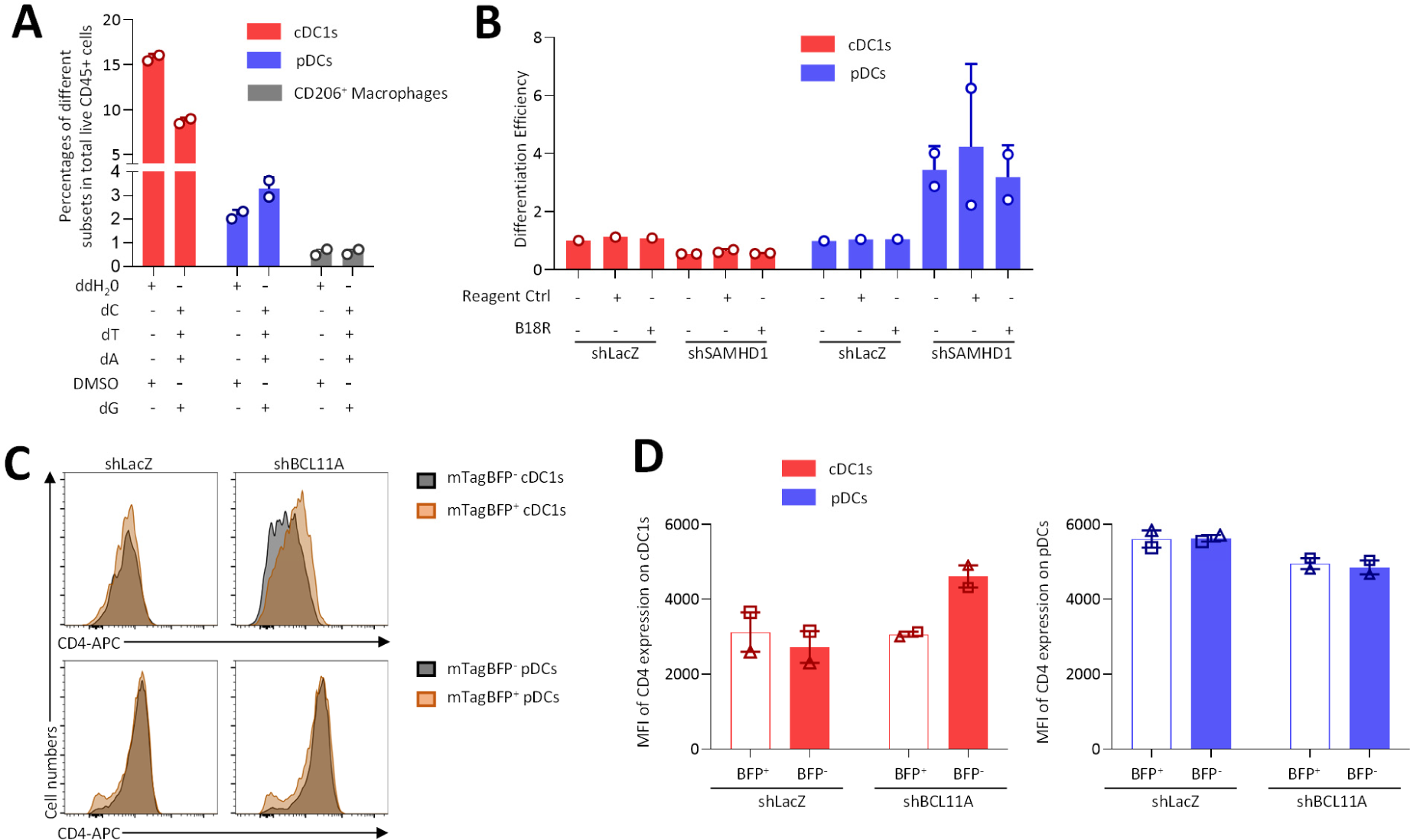
Investigation of putative mechanisms for SAMHD1 control of pDC/cDC differentiation and of BCL11A restriction activity against HIV-1 in cDC1s. (**A**) Impact of activation of the dN salvage pathway on the development of cDC1s, pDCs and macrophages. The results shown are from two independent experiments. (**B**) Impact of blockade of the IFN-I signaling pathway on the effects of *SAMHD1* knock-down on cDC1 and pDC development. The results shown are from two independent experiments for the shSAMHD1 condition. (**C**) Impact of *BCL11A* knock-down on CD4 expression levels on cDC1s and pDCs, as detected by FACS. The results shown are from one experiment representative of three independent ones. (**D**) CD4 MFI on cDC1s and pDCs upon *BCL11A* knock-down as compared to control lentivector transduction. The results are shown from two independent experiments, each with a different CB donor, with one shRNA.

